# Nanomedicines Targeting Signaling of Protease-activated Receptor 2 in Organelles Provide Sustained Analgesia

**DOI:** 10.1101/2024.09.10.612022

**Authors:** Shavonne Teng, Rocco Latorre, Divya Bhansali, Parker K. Lewis, Rachel E. Pollard, Chloe J. Peach, Badr Sokrat, Gokul SA Thanigai, Tracy Chiu, Dane D. Jensen, Nestor N Jimenez-Vargas, Abby Mocherniak, Lucas T Parreiras-E-Silva, Michel Bouvier, Matthew Bogyo, Michael M. Gaspari, Stephen J. Vanner, Nathalie M. Pinkerton, Kam Leong, Brian L. Schmidt, Nigel W. Bunnett

## Abstract

Although many internalized G protein-coupled receptors (GPCRs) continue to signal, the mechanisms and outcomes of GPCR signaling in organelles are uncertain due to the challenges of measuring organelle-specific signals and of selectively antagonizing receptors in intracellular compartments. Herein, genetically-encoded biosensors targeted to subcellular compartments were used to analyze organelle-specific signaling of protease-activated receptor 2 (PAR_2_); the propensity of nanoparticles (NPs) to accumulate in endosomes was leveraged to selectively antagonize intracellular PAR_2_ signaling of pain. PAR_2_ agonists evoked sustained activation of PAR_2_, Gαq and β-arrestin-1 in early, late and recycling endosomes and the *cis*- and *trans*-Golgi apparatus, and activated extracellular signal regulated kinase (ERK) in the cytosol and nucleus, measured with organelle-targeted biosensors. Dendrimer and core-shell polymeric NPs accumulated in early and late endosomes of HEK293 cells, colonic epithelial cells and nociceptors, detected by confocal imaging of fluorescent NPs. NPs efficiently encapsulated and slowly released AZ3451, a negative allosteric PAR_2_ antagonist. NP-encapsulated AZ3451, but not unencapsulated AZ3451, rapidly and completely reversed PAR_2_, Gαq and β-arrestin-1 activation in endosomes and the Golgi apparatus and ERK activation in the cytosol and nucleus. When administered into the mouse colon lumen, dendrimer NPs accumulated in endosomes of colonocytes and polymeric NPs targeted neurons, sites of PAR_2_ expression. Both NP-AZ3451 formulations, but not unencapsulated AZ3451, caused long-lasting analgesia and normalized aberrant behavior in preclinical models of inflammatory bowel disease. Thus, organelle-specific PAR_2_ signals in colonocytes and nociceptors mediate pain. Antagonism of PAR_2_ in organelles, rather than at the plasma membrane, provides effective pain relief.

**Significance Statement:** Once activated at the cell surface, many GPCRs internalize and continue to signal. The mechanisms and physiological relevance of intracellular GPCR signaling are uncertain. By using organelle-targeted biosensors, we detected sustained activation of the GPCR, PAR_2_, and its effectors in early, late and recycling endosomes, the *cis*- and *trans*-Golgi apparatus, and the cytosol and nucleus. NPs that delivered AZ3451, a PAR_2_ antagonist, to endosomes disrupted these intracellular signals, whereas unencapsulated AZ3451 was minimally effective. After intracolonic administration to mice, NPs accumulated in colonocytes and neurons. NP-encapsulated AZ3451, but not unencapsulated AZ3451, reversed pain in preclinical models of inflammatory bowel disease. Thus, intracellular PAR_2_ signaling mediates pain and antagonism of intracellular rather than plasma membrane PAR_2_ provides effective therapy.

## Introduction

G protein-coupled receptors (GPCRs) are a large family of seven transmembrane domain proteins that control many physiological and pathological processes (1). GPCRs are the single largest drug target, illustrating their importance in disease and medicine (2). GPCRs are conventionally considered as plasma membrane receptors that bind to agonists in the extracellular fluid and couple to heterotrimeric G proteins, which convey intracellular signals. Consequently, drug discovery has focused on targeting GPCRs at the plasma membrane. However, GPCR activity at the plasma membrane is usually transient. G protein-coupled receptor kinases phosphorylate activated GPCRs, which increases their affinity for β-arrestins (βARRs) (3). βARRs uncouple GPCRs from G proteins, desensitizing plasma membrane signaling, and couple GPCRs to clathrin and adaptor protein-2, leading to receptor endocytosis. Although endocytosis was formerly viewed as another mechanism to arrest signaling, many internalized GPCRs continue to signal, including those that form high affinity, sustained βARR interactions and some receptors that interact with βARRs with low affinity and transiently (4–13). GPCR signaling in endosomes mediates the actions of hormones (4, 6) and contributes to pain (8, 10, 13, 14). However, our understanding of the mechanisms and functions of intracellular GPCR signaling is hampered by the challenges of measuring GPCR and effector activity in specific organelles and of selectively antagonizing intracellular rather than cell-surface GPCRs. Moreover, whether antagonists disrupt preexisting GPCR signals in organelles, which requires penetration of plasma and organelle membranes, and engagement with GPCR conformations within multi-protein signaling complexes of acidic organelles, is seldom investigated. The inability of drugs to inhibit GPCR signals in organelles could contribute to their failure in clinical trials for the treatment of chronic diseases, where GPCRs are likely internalized due to over-production of agonists.

Herein, we measured the activity of GPCRs, Gα isoforms, βARRs and kinases in subcellular compartments by bioluminescence resonance energy transfer (BRET) and Förster response energy transfer (FRET) assays. BRET and FRET biosensors were targeted to the plasma membrane and organelles, which enabled analysis of compartmentalized signaling in live cells with high spatial and temporal resolution. We used nanoparticles (NPs) to encapsulate and deliver a GPCR antagonist to endosomes, which allowed determination of the contribution of intracellular GPCR signaling to pathology. While the mechanisms by which NPs traffic to endosomes are not fully understood, endocytosis of small NPs (*e.g.*, < 200 nm for clathrin-mediated endocytosis) is ubiquitous in non-phagocytic cells (15).

We investigated the mechanisms by which protease-activated receptor-2 (PAR_2_) signals in subcellular compartments and devised an approach to antagonize PAR_2_ signaling in organelles of key cell types for the treatment of pain. PAR_2_ is highly expressed in colonocytes and associated neurons and has been implicated in inflammatory bowel disease (IBD) and irritable bowel syndrome (IBS) (16). During inflammatory diseases involving the colon, mammalian and bacterial proteases cleave and activate PAR_2_, leading to receptor endocytosis and intracellular signaling (10, 14, 17). Since PAR_2_ signals from endosomes of colonocytes and nociceptors to evoke inflammation and pain (10, 14, 17), drug delivery systems must bypass the plasma membrane and deliver antagonists to endosomes to be effective. Accordingly, we asked the question whether a hydrophobic allosteric antagonist of PAR_2_, AZ3451 (18), loaded into NPs could be used to antagonize endosomal PAR_2_ and improve analgesic efficacy in preclinical mouse models of IBD. We studied two translatable but fundamentally different NP types: slow-release dendrimers (US NCT05205161) and core-shell polymeric NPs (US NCT03217838). We administered the NPs locally in the colon to mitigate the challenges and side effects of systemic administration and studied whether the NPs disrupt endosomal GPCR signaling and produce analgesic efficacy (19).

## Results

### PAR_2_ signals at the plasma membrane, in early, late and recycling endosomes, and in the *cis*- and *trans*-Golgi apparatus

PAR_2_, Gα and βARR signaling at the plasma membrane and in organelles was measured using BRET proximity assays with genetically-encoded biosensors targeted to subcellular microdomains of HEK293T cells, which are commonly used to study GPCR signaling. Enhanced bystander BRET (ebBRET), which exploits the affinity of pairs of *Renilla* (R)-tagged proteins (20, 21), was used to determine the kinetics of PAR_2_-induced trafficking of mini (m) Gαq, mGαi and βARR1 to organelles (**Fig. S1A**). mGα proteins are N-terminally truncated forms of Gα that interact with activated GPCRs (22, 23). Rluc8-mGαq, Rluc8-mGαi or βARR1-RlucII and a plasma membrane marker (CAAX) coupled to RGFP or markers of early endosomes (Rab5a), late endosomes (Rab7a), recycling endosomes (Rab4a, Rab11a), or the *cis*-Golgi network (Giantin) coupled to tandem (td)RGFP were coexpressed in HEK293T cells. The PAR_2_ agonists 2-furoyl-LIGRLO-NH_2_ (2F) (10 µM) and trypsin (100 nM) rapidly increased ebBRET between Rluc8-mGαq, Rluc8-mGαi or βARR1-RlucII and RGFP-CAAX (plasma membrane), tdRGFP-Rab5a (early endosomes), tdRGFP-Rab7a (late endosomes), tdRGFP-Rab4a and tdRGFP-Rab11a (recycling endosomes), and tdRGFP-Giantin (*cis*-Golgi apparatus) (**Fig. S1B-G**). mGαq and mGαi were recruited to the plasma membrane, endosomes and the Golgi apparatus with similar kinetics, although the recruitment of βARR1 to endosomes lagged behind its recruitment to the plasma membrane. ebBRET signals at the plasma membrane declined during the experiment, whereas most other signals were maintained for 20 min.

EbBRET is limited to studying the proximity of only two proteins. To overcome this limitation, NanoLuc Binary Technology BRET (NanoBiT-BRET, nbBRET) was used to simultaneously measure the proximity between PAR_2_, an effector (mGα or βARR), and a plasma membrane or organelle localization marker (14, 24) (**Fig. 1A**). NbBRET is based on a split luciferase assay whereby human PAR_2_ is C-terminally tagged with Natural Peptide (NatP) NanoLuc fragment (13 residue NanoBiT) and HA-tagged localization markers of the plasma membrane (CAAX), early endosomes (FYVE), late endosomes (Rab7a), recycling endosomes (Rab4a, Rab11a), *cis*-Golgi network (Giantin) and *trans*-Golgi network (TGN38) are tagged with large NanoBiT fragment (LgBiT). Luminescence occurs from a complex between PAR_2_-NatP and LgBiT markers, which serves as an energy donor for Venus-mGα or βARR-YFP. The expected localization of markers was confirmed (**Fig. S2**). NbBRET has been verified (14). 2F and trypsin rapidly increased nbBRET between PAR_2_-NatP, Venus-mGαq, Venus-mGαi or βARR1-YFP, and LgBiT-CAXX, LgBiT-FYVE, LgBiT-Rab7a, LgBiT-Rab4a, LgBiT-Rab11a, LgBiT-Giantin and TGN38-LgBiT in HEK293T cells (**Fig. 1B-G**). Responses at the plasma membrane and in endosomes were of similar kinetics and sustained for 20 min, although signals in the Golgi apparatus were delayed. 2F and trypsin did not affect nbBRET between PAR_2_-NatP, Venus-mGαs and any organelle marker (**Fig. 1H, I**).

**Figure 1.**
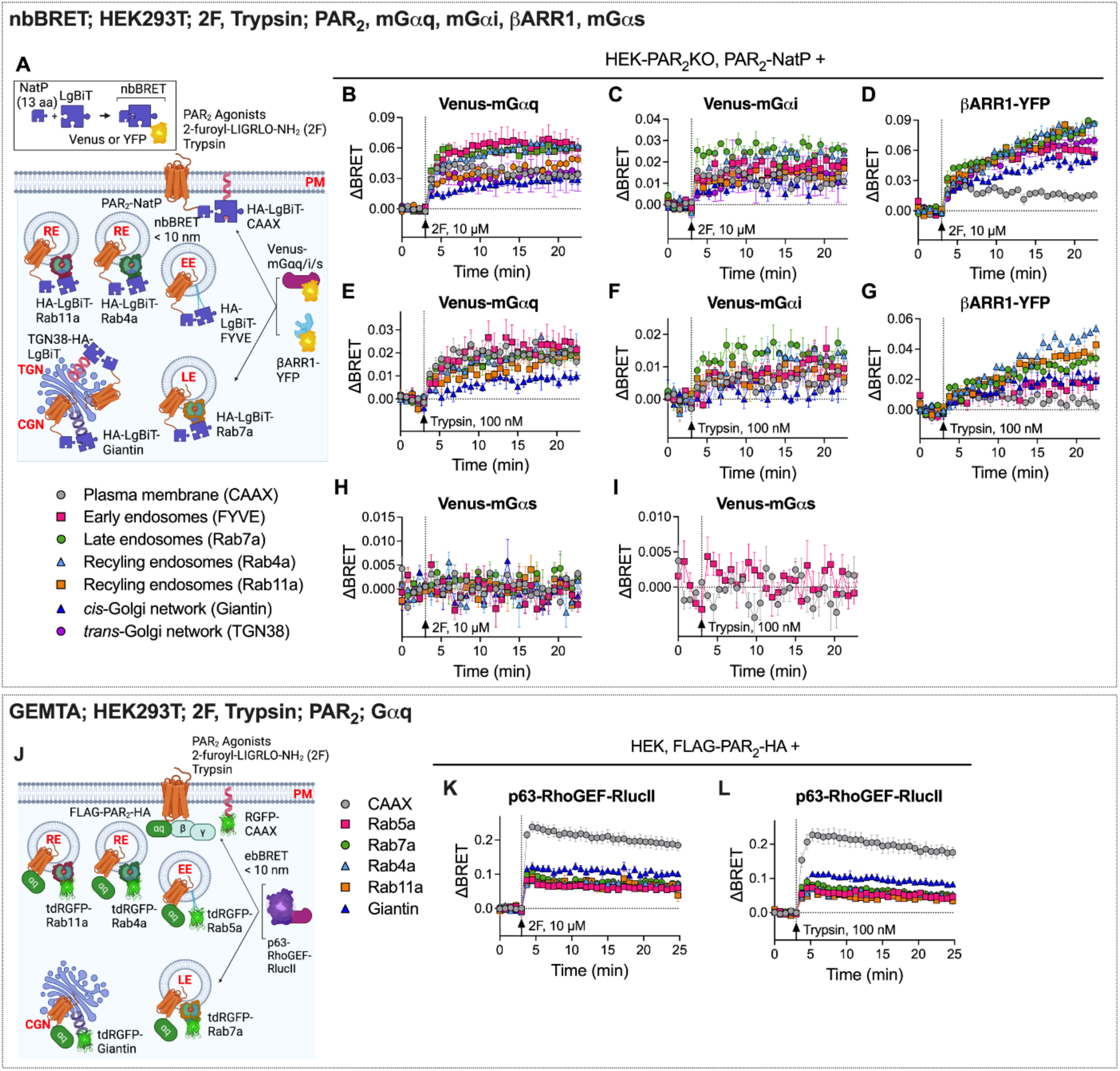
PAR_2_ signals from the plasma membrane, endosomes, and the Golgi apparatus. **A**. NbBRET assay of recruitment of mGαq (**B**, **E**), mGαi (**C, F**), mGαs (**H, I**) βARR1 (**D**, **G**) of Gαi (**H, I**) to PAR_2_ to the plasma membrane (CAAX), early endosomes (FYVE), late endosomes (Rab7a), recycling endosomes (Rab4a, Rab11a), *cis*-Golgi network (CGN) or *trans*-Golgi network (TGN) of HEK293T cells. **J**. GEMTA assays of PAR_2_ and Gαq signaling in HEK293T cells showing 2F-induced (**K**) or trypsin-induced (**L**) recruitment of p63-RhoGEF to the plasma membrane, early endosomes, late endosomes, recycling endosomes, and *cis*-Golgi-network. Mean±SEM, triplicate observations, n=5 experiments.

An ebBRET-based G protein effector membrane translocation assay (GEMTA) was used to monitor PAR_2_-induced activation of Gαq in subcellular compartments (25) (**Fig. 1J**). HEK293SL cells were transfected with FLAG-PAR_2_-HA, p63-RhoGEF-RlucII and plasma membrane marker CAAX fused to RGFP or intracellular markers (Rab5a, Rab7a, Rab4a, Rab11a, Giantin) fused to tdRGFP. 2F or trypsin rapidly stimulated ebBRET between p63-RhoGEF-RlucII and RGFP-tagged markers of the plasma membrane, early, late and recycling endosomes, and the *cis*-Golgi apparatus with similar kinetics (**Fig. 1K, L**). Signals were maintained during the period of observation (20 min).

The combined results provide evidence that the PAR_2_ agonists 2F and trypsin activate PAR_2_, Gαq, Gαi and βARR1 at the plasma membrane, in early, late and recycling endosomes, and in the *cis*- and *trans*-Golgi apparatus of HEK293 cells. These signals are sustained for at least 20 min.

### Dendrimer and core-shell polymeric NPs efficiently encapsulate and slowly release a PAR_2_ antagonist

By leveraging the prevalent endosomal uptake of NPs and the acidity of the endosomal lumen to trigger NP disassembly and cargo release, we can preferentially deliver GPCR antagonists to the desired site of action within endosomes. We have previously encapsulated antagonists of the neurokinin 1 receptor (NK_1_R) and calcitonin-like receptor (CLR) into soft polymer NPs designed to undergo endocytosis and disassemble and release antagonist cargo in acidic endosomes (26–28). After intrathecal or periorbital injection, pH-responsive NPs that deliver NK_1_R and CLR antagonists to endosomes of spinal neurons and periorbital Schwann cells, respectively, provide sustained antinociception in preclinical models of inflammatory, neuropathic and migraine pain (26–28). However, these routes of administration are challenging in a clinical setting. Moreover, pH-responsive NPs are unsuitable for antagonism of receptors at peripheral sites of inflammation and injury because they would prematurely disassemble in the acidified extracellular environment of diseased tissues (29) rather than in endosomes. While immediate pH-dependent antagonist release from NPs is ideal for the rapid reversal of acute pain (26–28), sustained release of antagonists is optimal for long-lasting alleviation of chronic pain. To antagonize intracellular PAR_2_ in inflamed tissues and provide long-lasting analgesia, dendrimer and core-shell polymeric NPs were developed for sustained and pH-independent delivery of AZ3451, a hydrophobic negative allosteric antagonist of PAR_2_ (18), to endosomes.

Poly(amidoamine) PAMAM dendrimers self-assemble into NPs with defined size, shape and a hydrophobic core that can encapsulate hydrophobic drug candidates (30, 31). Cholesterol (Chol) chloroformate was used to modify the surface of amine-terminated generation 3 PAMAM-dendrimers, forming soft, cationic PAMAM-Chol NPs with mucoadhesive properties (32, 33). AZ3451 was entrapped in the core during self-assembly of the cholesterol-functionalized PAMAM-G3 in aqueous solution, forming PAMAM-Chol-AZ (**Fig. 2A**). AZ3451 was encapsulated with 99.6% efficiency and 20% (w/w) loading. PAMAM-Chol-AZ had a hydrodynamic diameter of 223±3 nm (mean±SD), a polydispersity index (PDI) <0.2 and a positive zeta potential of +65±1 mV (**Fig. 2B**). Empty PAMAM-Chol-Ø had a hydrodynamic diameter of 129±5 nm and a zeta potential of +53±1 mV. PAMAM-Chol-AZ NPs were unaggregated and uniformly spherical (**Fig. 2C**). Passive release of AZ3451 from the dendrimer was observed for at least 48 h (**Fig. 2D)**. AZ3451 was released at a similar rate at pH 5.5, 6.5 and 7.4, representing endosomal and extracellular environments. Fluorescent cyanine 5 (Cy5) was conjugated to PAMAM-Chol using N-hydroxysuccinimide (*NHS*) chemistry. PAMAM-Chol-Cy5 was of similar size and charge to its unlabeled counterpart (**Fig. 2B**). PAMAM-Chol-Ø (0.3-10 µg/mL) did not cause cytotoxicity of HEK293T cells after 4 or 24 h (**Fig. 2E**). The highest concentration (10 µg/mL) corresponds to 3.3 µM PAPM-Chol-AZ, which exceeds the maximum concentration tested in signaling assays.

**Figure 2.**
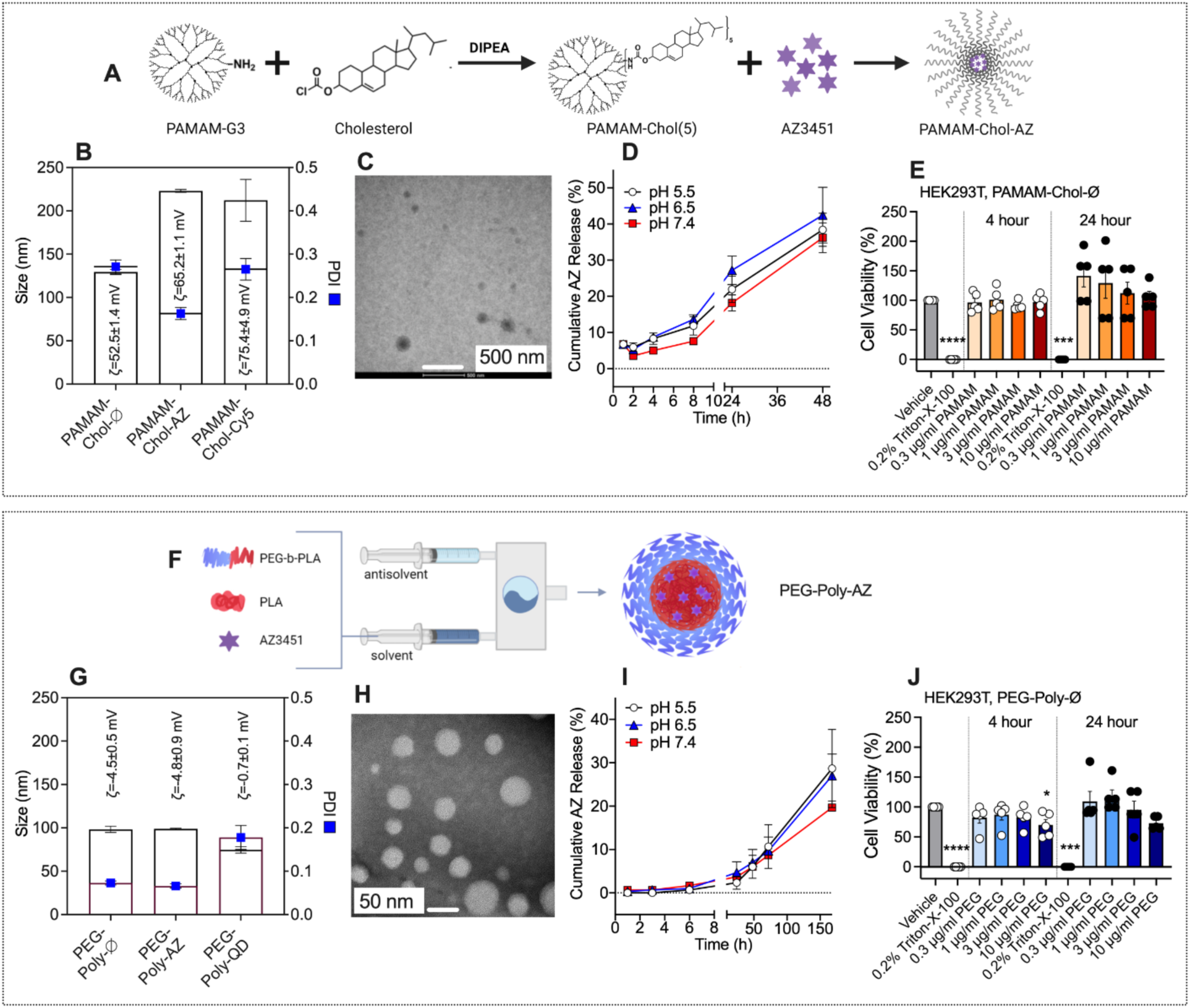
NPs efficiently encapsulate and slowly release a PAR_2_ antagonist. **A-E.** PAMAM-Chol NPs. **F-J.** PEG-Poly NPs. **A, F.** Composition and assembly of NPs. **B, G.** Size and PDI of NPs. **C, H.** TEM images of NPs. **D, I.** Kinetics of AZ3451 release from NPs *in vitro* at pH 5.5, 6.5 and 7.4. **B, G, D, I.** Mean±SEM, triplicate observations, n=3 independent experiments. **C, H.** Representative images, n=3 independent experiments. **E, J.** Viability of HEK293T cells incubated with vehicle, Triton-X-100 (positive control) or graded concentrations of PAMAM-Chol-Ø or PEG-Poly-Ø NPs for 4 h or 24 h. Mean±SEM, triplicate observations, n=5 independent experiments. \**P*<0.05. One-way ANOVA, Dunnett’s multiple comparisons compared to vehicle.

Flash Nanoprecipitation (FNP) permits assembly of core-shell polymeric NPs by diffusion-limited self-assembly driven by hydrophobic interactions (34, 35). FDA approved poly(lactic acid)-poly(ethylene glycol) (PEG-*b*-PLA) was selected as the stabilizing block copolymer to create a biocompatible NP with a neutral, non-fouling surface. PEG was chosen to achieve a neutral outer PEG brush layer, known to enable deep tissue penetration (36), whereas PLA was selected as the hydrophobic block for its biodegradability and glassy morphology (37). An organic stream of tetrahydrofuran (THF) containing dissolved AZ3451, PLA and PEG-PLA was rapidly mixed against ultra-pure water streams to form core-shell PEGylated polymeric-AZ3451 (PEG-Poly-AZ) (**Fig. 2F**). AZ3451 encapsulation efficiency was 99.1%, which resulted in NP with 8% (w/w) AZ3451 loading. PEG-Poly-AZ had a hydrodynamic diameter of 99±1 nm, a PDI of <0.07 and a near neutral zeta potential of -5±1 mV (**Fig. 2G**). Empty PEG-Poly-Ø had an average hydrodynamic diameter of 98±6 nm and a zeta potential of -5±1 mV. PEG-Poly-AZ NPs were unaggregated and uniformly spherical (**Fig. 2H**). The *in vitro* release profiles of AZ3451 at pH 5.5, 6.5 and 7.4 were similar, demonstrating sustained release for at least 160 h (**Fig. 2I)**. PEG-Poly encapsulating fluorescent quantum dots (QD) had a PDI of <0.2, a hydrodynamic diameter of 99±1 nm and a zeta potential of -0.7±0.1 mV. PEG-Poly-Ø (0.3-10 µg/mL) did not induce toxicity after 4 or 24 h, with the exception of a slight decrease in viability at the highest concentration of PEG-Poly-Ø (**Fig. 2J**).

### Dendrimer and core-shell polymeric NPs accumulate in early and late endosomes

NP uptake was studied in HEK293T cells coexpressing Rab5a-RFP (early endosomes) and Rab7a-GFP (late endosomes). PAMAM-Chol-Cy5 and PEG-Poly-QD were detected in early and late endosomes after 2 and 4 h (**Fig. 3A-F**). The proportion of early and late endosomes containing PAMAM-Chol-Cy5 (2 h, early endosomes 92%; late endosomes 88%) exceeded the proportion of early and late endosomes containing PEG-Poly-QD (2 h, early endosomes 23%; late endosomes 31%). Thus, both NP formulations can be used to deliver cargo to endosomes.

**Figure 3.**
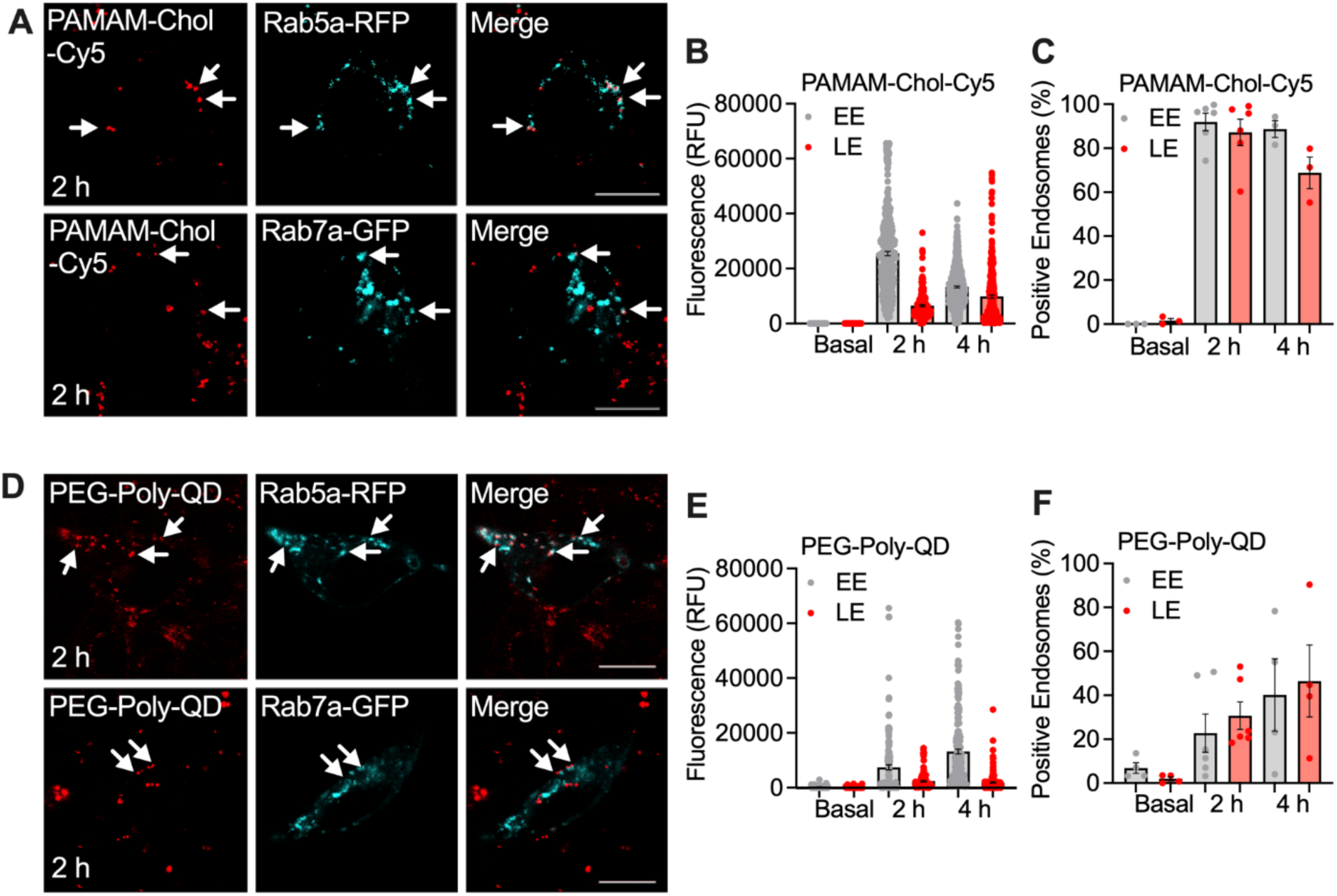
NPs accumulate in early and late endosomes. Uptake of PAMAM-Chol-Cy5 NPs (**A-C**) and PEG-Poly-QD NPs (**D-F**) in HEK293T cells. **A, D.** Localization of PAMAM-Chol-Cy5 NPs (**A**) and PEG-Poly-QD NPs (**D**) in early endosomes containing Rab5a-RFP and late endosomes containing Rab7a-GFP after 2 h. **B, E.** Quantification of PAMAM-Chol-Cy5 (**B**) and PEG-Poly-QD (**E**) fluorescence (relative fluorescence units, RFU) into early endosomes (EE) and late endosomes (LE). **C, F.** Proportion of early and late endosomes containing PAMAM-Chol-Cy5 (**C**) or PEG-Poly-QD (**F**) NPs. **A, D.** Representative images, n=4-6 independent experiments. **B, C, E, F.** Mean±SEM, triplicate observations, n=4-6 independent experiments.

### NP encapsulation enhances antagonism of PAR_2_ signaling in early, late and recycling endosomes, and in the *cis*- and *trans*-Golgi apparatus

Inflammation of the colon induces PAR_2_ redistribution from the plasma membrane to endosomes of colonocytes and nociceptors, which is attributable to activation of proteases in inflamed tissues (10, 14). Persistent endosomal signaling of PAR_2_ in colonocytes induces redistribution of tight junctions and mucosal influx of proinflammatory macromolecules from the lumen (14, 17), and sustained endosomal PAR_2_ signaling in nociceptors causes hyperexcitability and pain (10). Suppression of ongoing inflammation and pain therefore requires that antagonists penetrate plasma and endosomal membranes and engage with GPCRs within multi-protein signaling complexes of acidic endosomes (24). The ability of antagonists to reverse ongoing signals is seldom tested.

To determine whether NP encapsulated and unencapsulated AZ3451 reverse ongoing activation of PAR_2_ at the plasma membrane and in organelles, nbBRET was measured between PAR_2_-NatP, Venus-mGαq or βARR1-YFP, and LgBiT-tagged markers of the plasma membrane, endosomes and Golgi apparatus in HEK293T cells. Cells were challenged with 2F or trypsin and then exposed to PAMAM-Chol-AZ, PEG-Poly-AZ, unencapsulated AZ3451 (1, 3 µM) or vehicle (control) 7 min later when nbBRET was stable (**Fig. 4A, Fig. S3A, Fig. S4A, Fig. S5A**). In vehicle-treated cells, 2F and trypsin stimulated nbBRET between PAR_2_-NatP, Venus-mGαq or βARR1-YFP, and markers of the plasma membrane, early, late and recycling endosomes, and the *cis*- and *trans*-Golgi apparatus that was stable for at least 30 min (**Fig. 4B-G, Fig. S3B-I, Fig. S4B-H, Fig. S5B-H**). Unencapsulated AZ3451 caused a small, delayed and usually insignificant inhibition. In contrast, PAMAM-Chol-AZ and PEG-Poly-AZ both caused a faster, larger and significant inhibition that was complete within 30 min. NP-encapsulated AZ3451 was particularly more effective than unencapsulated AZ3451 at reversing 2F-stimulated nbBRET between PAR_2_-NatP and Venus-mGαq in late endosomes and the Golgi apparatus and in reversing 2F-stimulated nbBRET between PAR_2_-NatP and βARR1-YFP in early, late and recycling endosomes and the Golgi apparatus. NP-encapsulated AZ3451 was markedly more effective than unencapsulated AZ3451 at reversing trypsin-stimulated nbBRET between PAR_2_-NatP and Venus-mGαq in the Golgi apparatus and in reversing trypsin-stimulated nbBRET between PAR_2_-NatP and βARR1-YFP in early, late and recycling endosomes and the Golgi apparatus.

**Figure 4.**
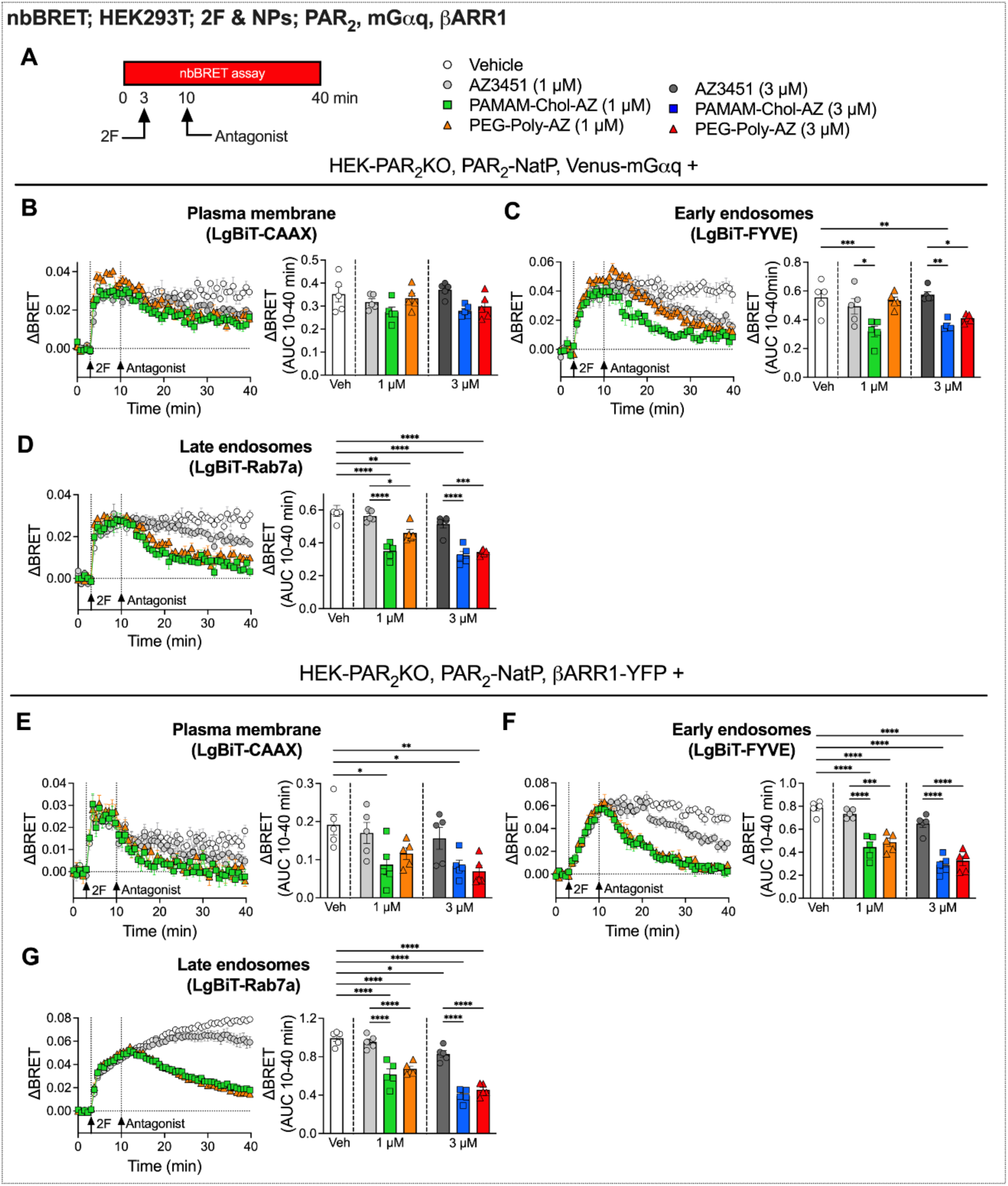
NPs encapsulating a PAR_2_ antagonist reverse 2F-stimulated PAR_2_ and βARR signaling in endosomes. **A**. NbBRET assays of PAR_2_ antagonist reversal of 2F-stimulated PAR_2_ and mGαq/βARR1 compartmentalized signaling in HEK293T cells. **B-D**. 2F-induced recruitment of mGαq to PAR_2_ to the plasma membrane (CAAX), early endosomes (FYVE), and late endosomes (Rab7a) in HEK293T cells followed by treatment with vehicle, unencapsulated AZ3451, PAMAM-Chol-AZ, or PEG-Poly-AZ (1 µM or 3 µM). Time courses show responses to 2F followed by vehicle or antagonist (1 µM). Area under curve (AUC, 10-40 min) show integrated responses to 2F after vehicle or antagonist (1 or 3 µM). **E-G.** 2F-induced recruitment of βARR1 to PAR_2_ to the plasma membrane (CAAX), early endosomes (FYVE), and late endosomes (Rab7a) in HEK293T cells followed by treatment of vehicle, unencapsulated AZ3451, PAMAM-Chol-AZ, or PEG-Poly-AZ (1 µM or 3 µM). Time courses show responses to 2F followed by vehicle or antagonist (1 µM). Area under curve (AUC, 10-40 min) show integrated responses to 2F after vehicle or antagonist (1 or 3 µM). Mean±SEM, triplicate observations, n=5 independent experiments. \**P*<0.05, \*\**P*<0.01, \*\*\**P*<0.001, \*\*\*\**P*<0.0001. One-way ANOVA, Šidák’s test.

GEMTA was used to monitor the effects of NPs on PAR_2_-induced activation of Gαq in subcellular compartments of HEK293SL cells coexpressing FLAG-PAR_2_-HA, p63-RhoGEF-RlucII and plasma membrane marker fused to RGFP or endosomal or Golgi markers fused to tdRGFP. Cells were challenged with 2F or trypsin and were exposed to PAMAM-Chol-AZ, PEG-Poly-AZ, unencapsulated AZ3451 or vehicle 2 min later when ebBRET was stable (**Fig. 5A, Fig. S6A, Fig. S7A**). In vehicle-treated cells, 2F and trypsin stimulated ebBRET between p63-RhoGEF-RlucII and RGFP-tagged markers of the plasma membrane, early, late and recycling endosomes, and the *cis*-Golgi apparatus that was stable for at least 20 min (**Fig. 5B-D, Fig. S6B-D, Fig. S7B-G**). Unencapsulated AZ3451 had a delayed, incomplete and usually insignificant inhibitory action, whereas PAMAM-Chol-AZ and PEG-Poly-AZ completely and significantly inhibited ebBRET at the plasma membrane, endosomes and the Golgi apparatus. Strikingly, unencapsulated AZ3451 did not significantly inhibit 2F-stimulated ebBRET between p63-RhoGEF-RlucII and RGFP-tagged markers of late or recycling endosomes or the Golgi apparatus and did not significantly inhibit trypsin-stimulated ebBRET between p63-RhoGEF-RlucII and RGFP-tagged markers of any subcellular compartment. In sharp contrast, PAMAM-Chol-AZ and PEG-Poly-AZ completely and significantly inhibited ebBRET signals in all compartments within ∼10 min.

**Figure 5.**
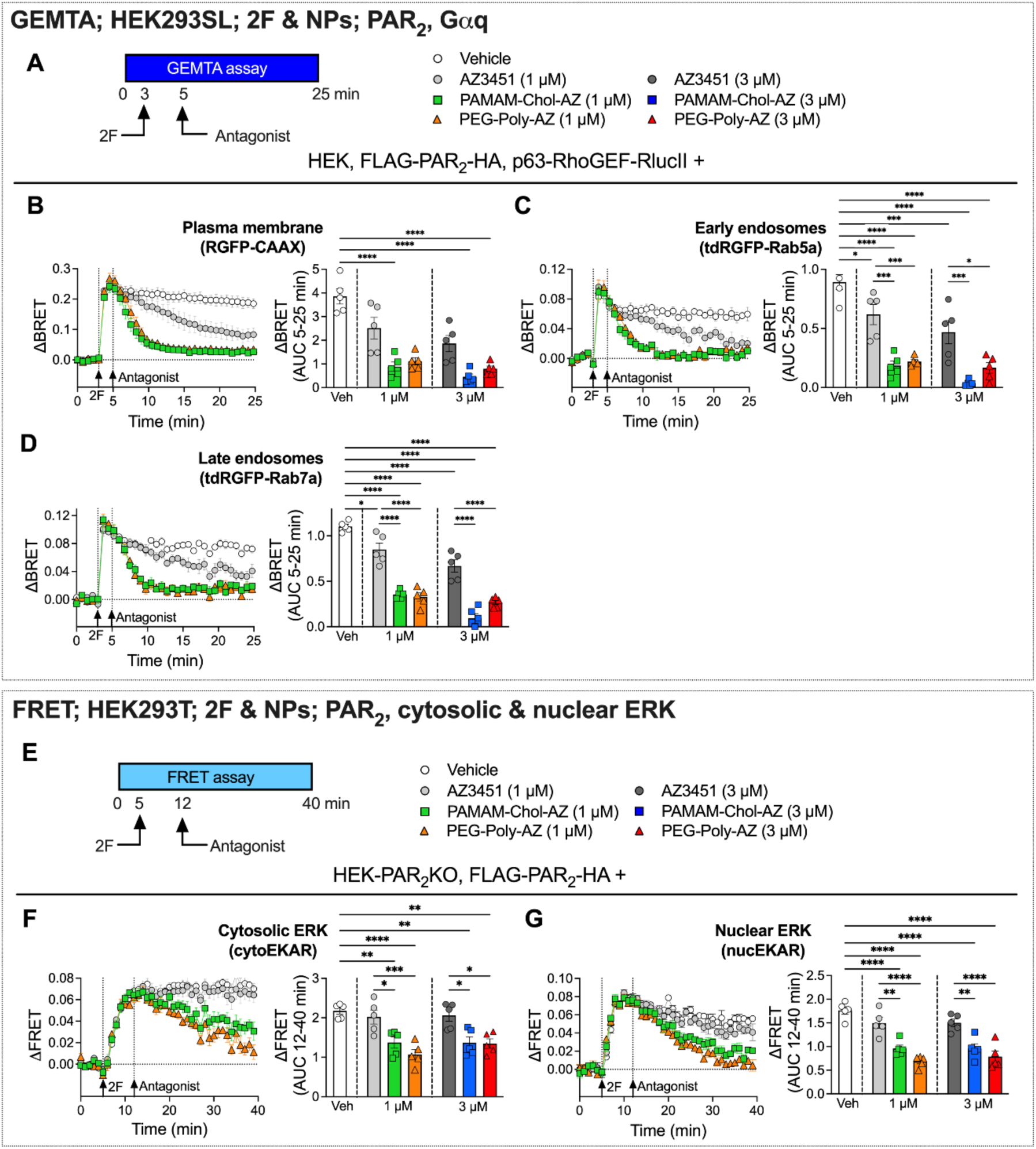
NPs encapsulating a PAR_2_ antagonist reverse 2F-stimulated Gαq signaling in endosomes and ERK signaling in the cytosol and nucleus. **A**. GEMTA assays of PAR_2_ antagonist reversal of 2F-stimulated PAR_2_ and Gαq compartmentalized signaling in HEK293T cells. **B-D**. 2F-induced recruitment of p63-RhoGEF to the plasma membrane (CAAX), early endosomes (Rab5a), and late endosomes (Rab7a) followed by treatment with vehicle, unencapsulated AZ3451, PAMAM-Chol-AZ, or PEG-Poly-AZ (1 µM or 3 µM). Time courses show responses to 2F followed by vehicle or antagonist (1 µM). Area under curve (AUC, 5-25 min) show integrated responses to 2F after vehicle or antagonist (1 or 3 µM). **E**. FRET assays of ERK activity in HEK293T cells. **F,G**. 2F-induced activation of cytosolic (cytoEKAR, **F**) and nuclear (nucEKAR, **G**) ERK followed by treatment with vehicle or antagonists. Time courses show responses to 2F followed by vehicle or antagonists (1 µM). Area under curve (AUC, 12-40 min) show integrated responses to 2F after vehicle or antagonists (1 or 3 µM). Mean±SEM, triplicate observations, n=5 experiments. \**P*<0.05, \*\**P*<0.01, \*\*\**P*<0.001, \*\*\*\**P*<0.0001. One-way ANOVA, Šidák’s test.

PAR_2_ endosomal signaling activates ERK (5), which mediates the proinflammatory and algesic actions of PAR_2_ in the colon (10, 17). To determine whether NP-encapsulated AZ3451 could reverse compartmentalized ERK signaling, FRET biosensors targeted to the cytoplasm (cytoEKAR) or nucleus (nucEKAR) were expressed with FLAG-PAR_2_-HA in HEK293T cells. Cells were challenged with 2F and were then exposed to PAMAM-Chol-AZ, PEG-Poly-AZ, unencapsulated AZ3451 or vehicle 7 min later when responses were stable (**Fig. 5E**). 2F-stimulated ERK activity in the cytoplasm and nucleus was sustained for 35 min in vehicle-treated cells (**Fig. 5F, G**). Unencapsulated AZ3451 was unable to inhibit cytosolic or nuclear ERK, whereas PAMAM-Chol-AZ and PEG-Poly-AZ completely reversed ERK activity within 35 min.

To ascertain whether NP encapsulation affects the potency of AZ3451, HEK293T cells were preincubated with vehicle, PAMAM-Chol-AZ, PEG-Poly-AZ or unencapsulated AZ3451 for 4 h (to allow cell penetration and antagonist release) and were then challenged with 2F (**Fig. S8A**). PAMAM-Chol-AZ (**Fig. S8B**), PEG-Poly-AZ (**Fig. S8C**) and unencapsulated AZ3451 (**Fig. S8D**) concentration-dependently inhibited 2F-induced PAR_2_ and mGαq nbBRET in all subcellular compartments with similar potency and efficacy. Equivalent solids concentrations of PAMAM-Chol-Ø (**Fig. S8E**) or PEG-Poly-Ø (**Fig. S8F**) NPs had no effect. These results show that AZ3451 retains full antagonistic activity when encapsulated into NPs.

Thus, when encapsulated into two different NPs that accumulate in endosomes, AZ3451 effectively reverses activation of PAR_2_ and its Gαq and βARR1 effectors in endosomes and the Golgi apparatus. The downstream consequence is abrogation of ERK signaling in the cytosol and nucleus, which underlies the proinflammatory and pronociceptive functions of PAR_2_. Unencapsulated AZ3451 does not effectively suppress ongoing signals. Empty NPs are inactive.

### NP-encapsulated antagonist blocks PAR_2_ actions on colonic epithelial cells and nociceptors

The ability of NP-encapsulated AZ3451 to antagonize PAR_2_ was studied in colonic epithelial cells and dorsal root ganglion (DRG) neurons that endogenously express PAR_2_ and mediate inflammation and pain.

Endosomal signaling of PAR_2_ in T84 colonic epithelial cells induces the redistribution of tight junctional proteins, which increases paracellular permeability and causes inflammation (14, 17). PAMAM-Chol-Cy5 and PEG-Poly-QD accumulated in early endosomes of T84 cells within 60 min, identified by early endosomal antigen 1 (EEA1) immunostaining (**Fig. 6A, B**). The PAR_2_ agonist 2F (100 µM) increased nbBRET between PAR_2_-NatP, Venus-mGαq or βARR1-YFP, and LgBiT-CAXX, LgBiT-FYVE, LgBiT-Rab7a, LgBiT-Rab4a and LgBiT-Rab11a in T84 cells (**Fig. 6C-E**). Preincubation with PAMAM-Chol-AZ or PEG-Poly-AZ (1 µM) but not with empty control PAMAM-Chol-Ø or PEG-Poly-Ø inhibited 2F-stimulated nbBRET between PAR_2_ and mGαq at the plasma membrane, early, late and recycling endosomes of T84 cells.

**Figure 6.**
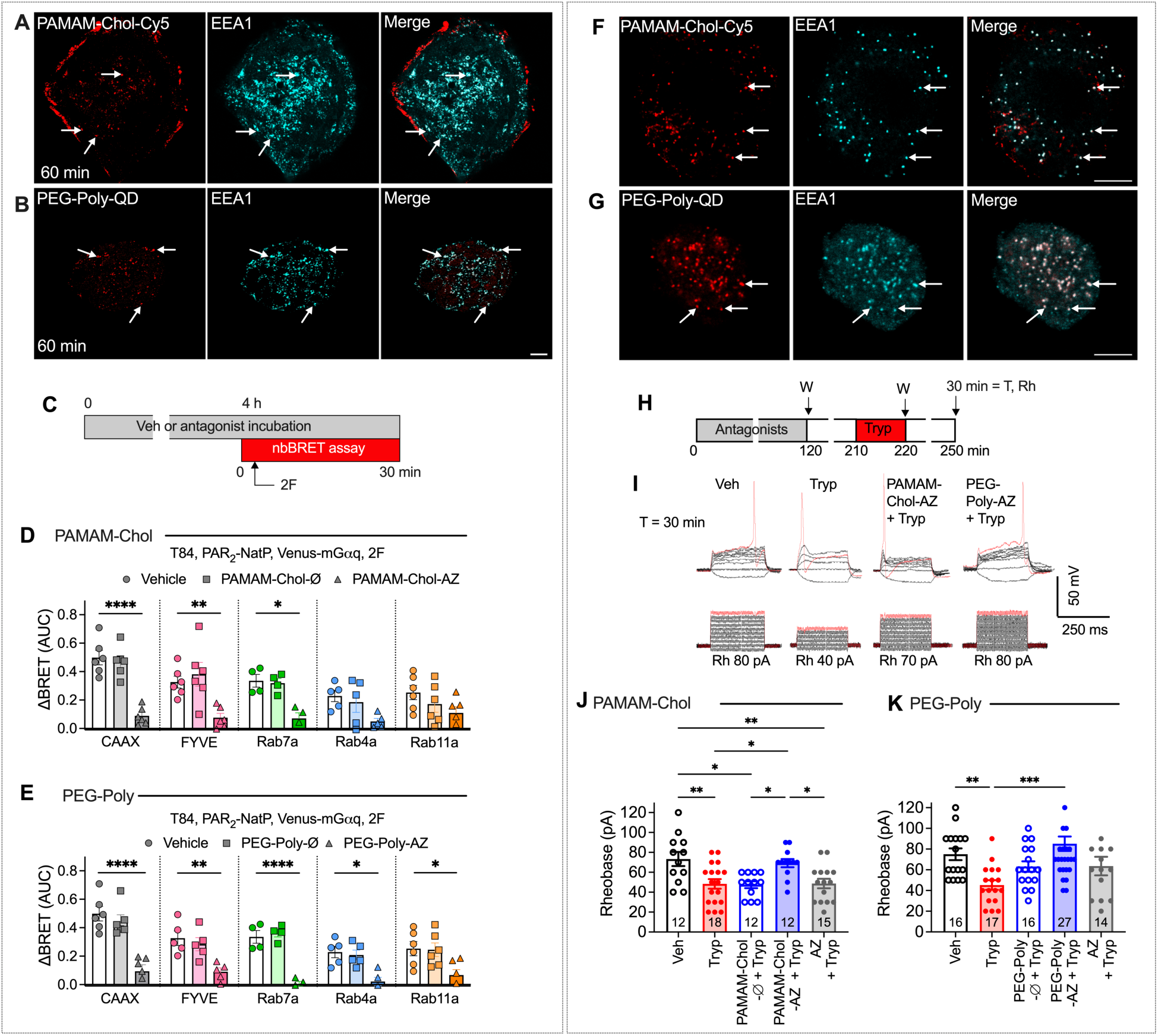
NPs that accumulate in endosomes of colonic epithelial cells and nociceptors cause sustained inhibition of PAR_2_-induced signaling and sensitization. **A-E.** T84 cells. **A, B.** PAMAM-Chol-Cy5 (**A**) and PEG-Poly-QD (**B**) uptake into early endosomes (EEA1+ve). Representative images, n=3 experiments. Scale, 20 µm. **C.** NbBRET assays of 2F-induced recruitment of mGαq to PAR_2_ at the plasma membrane, early endosomes, late endosomes and recycling endosomes of T84 cells preincubated with vehicle or PAMAM-Chol-AZ (1 µM), PAMAM-Chol-Ø (**D**) or PEG-Poly-AZ (1 µM) or PEG-Poly-Ø (**E**). Mean±SEM, triplicate observations, n=5 experiments. \**P*<0.05, \*\**P*<0.01, \*\*\**P*<0.001, \*\*\*\**P*<0.0001. One-way ANOVA, Šidák’s test. **F-J.** Mouse DRG neurons. **F, G.** PAMAM-Chol-Cy5 (**F**) and PEG-Poly-QD (**G**) uptake into early endosomes (EEA1+ve) of neurons. Representative images, n=5 experiments. Scale, 10 µm. **H-K.** Patch clamp recordings of rheobase (Rh). **H.** Neurons were untreated or incubated with antagonists for 120 min, washed (W) and recovered in antagonist-free medium for 90 min, challenged with trypsin (Tryp) for 10 min, washed, and rheobase measured at T = 30 min post-trypsin. **I.** Representative recordings. **J, K.** Rheobase after pretreatment with vehicle, or unencapsulated AZ3451, or PAMAM-Chol-Ø, PAMAM-Chol-AZ (**J**) or PEG-Poly-Ø, PEG-Poly-AZ (**K**). Mean±SEM, n, neurons number (in bars), 4-8 mice. \**P*<0.05, \*\**P*<0.01, \*\*\**P*<0.001. **J.** One-way ANOVA, Tukey’s Test. **K.** Kruskal-Wallis, Dunn’s test.

Endosomal signaling of PAR_2_ in DRG nociceptors induces sustained sensitization, consistent with pain (10). PAMAM-Chol-Cy5 and PEG-Poly-QD accumulated in EEA1+ve early endosomes of NeuN+ve neurons in cultures of mouse DRG after 60 min (**Fig. 6F, G**). To determine whether NP-encapsulated and unencapsulated AZ3451 inhibit PAR_2_-induced sensitization, neurons were untreated or incubated with PAMAM-Chol-Ø, PAMAM-Chol-AZ, PEG-Poly-Ø, PEG-Poly-AZ or unencapsulated AZ3451 (120 min, 100 nM AZ3451); neurons were washed, recovered in antagonist-free medium for 90 min, challenged with trypsin (50 nM, 10 min), and washed again. Rheobase (minimum input current to fire an action potential) was measured at T=30 min post-trypsin by perforated-patch clamp recording (**Fig. 6H**). Trypsin decreased the rheobase to 48.33±4.8 pA (34.1±6.5% decrease, *P*<0.01) (**Fig. 6J**) and 45.29±4.6 pA (39.6±6.1% decrease, *P*<0.01) (**Fig. 6K**) compared to vehicle (rheobase 73.33±7.1 pA and 75±5.7 pA). PAMAM-Chol-Ø, PEG-Poly-Ø and unencapsulated AZ3451 did not affect the response to trypsin, whereas PAMAM-Chol-AZ and PEG-Poly-AZ prevented trypsin-induced hyperexcitability of nociceptors (rheobase 69.16±4.1 pA and 85.18±6.9 pA respectively).

Thus, NP-encapsulated AZ3451 blocks intracellular PAR_2_ signals in colonic epithelial cells and causes a sustained and complete inhibition of PAR_2_-evoked hyperexcitability of DRG neurons.

### Luminally administered NPs accumulate in endosomes of epithelial cells and neurons in the colon

Observations of knockin mice expressing PAR_2_ C-terminally fused to monomeric ultrastable GFP (muGFP) reveal that PAR_2_ redistributes from the plasma membrane to endosomes in the inflamed colon (14), which is attributable to the activation of proteases from immune cells (38). Since PAR_2_ signals from endosomes of colonocytes and neurons to evoke inflammation and pain, the delivery and endosomal retention of NP-encapsulated antagonists to both cell types is therapeutically desirable.

To examine cellular and organelle targeting of NPs, PAMAM-Chol-Cy5 and PEG-Poly-QD were administered into the colon lumen by enema, which circumvented the challenges of gastric acidification and small intestinal transit associated with oral delivery. NP uptake was examined by confocal imaging of excised colon. PAMAM-Chol-Cy5 was prominently detected in colonocytes at the tips of mucosal folds within 1 h, with maximal signals at 2 h and retention for 8 h (**Fig. 7A**). Conversely, PEG-Poly-QD was rarely detected in colonocytes but instead penetrated into the lamina propria, submucosa and musculature (**Fig. 7A**). Quantification of fluorescent signals revealed preferential targeting of PAMAM-Chol-Cy5 to the mucosa, consistent with colonocyte uptake, and of PEG-Poly-QD to the muscle layer, consistent with accumulation in the myenteric plexus (**Fig. 7B**), and confirmed retention of NPs in tissues for at least 8 h.

**Figure 7.**
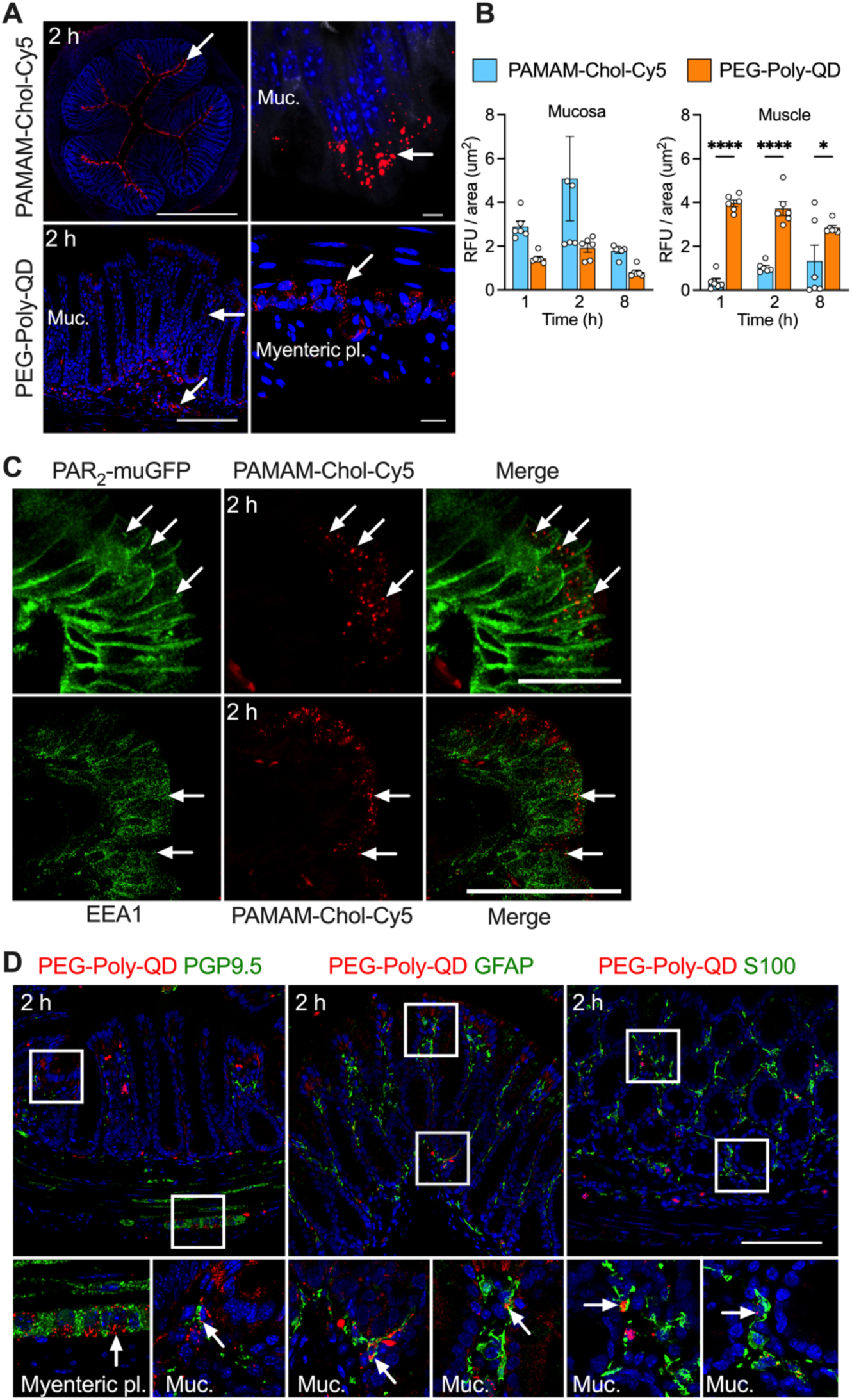
Luminally administered PAMAM-Chol-Cy5 and PEG-Poly-QD NPs accumulate in endosomes of colonocytes, neurons and glial cells. **A.** Uptake of luminally administered PAMAM-Chol-Cy5 into colonocytes and PEG-Poly-QD into enteric neurons after 2 h. Muc., mucosa; myenteric pl., myenteric plexus. **B.** Time course of uptake of PAMAM-Chol-Cy5 and PEG-Poly-QD into the mucosa and muscle layers of the colon. Mean±SEM, n=5 mice. \**P*<0.05, \*\*\*\**P*<0.0001. T-test. **C.** Localization of PAMAM-Chol-Cy5 in colonocytes expressing PAR_2_-muGFP and detection of PAMAM-Chol-Cy5 in EEA-1+ve early endosomes after 2 h. Arrows indicate PAMAM-Chol-Cy5 localization in endosomes containing PAR_2_-muGFP and EEA1. **D.** Localization of PEG-Poly-QD in the lamina propria of the mucosa and association with PGP9.5+ve neurons and GFAP+ve and S100+ve glial cells in the mucosa and muscle layers of the colon after 2 h. Lower images correspond to white boxes in upper images. Representative images, 4 (**A**) and 3 (**C, D**) experiments. Scale, 100 µm.

Targeting of NPs to specific cells and organelles was examined by immunostaining. When administered into the colon lumen of *Par_2_-mugfp* mice, PAMAM-Chol-Cy5 was detected in colonocytes expressing PAR_2_-muGFP and sometimes colocalized with PAR_2_-muGFP in endosomes (**Fig. 7C**). PAMAM-Chol-Cy5 colocalized with EEA1 in endosomes of colonocytes (**Fig. 7C**). In contrast, PEG-Poly-QD accumulated in cells of the nervous system in the mucosa, submucosa and muscle layers of the colon. PEG-Poly-QD was detected in neurons, identified by immunostaining for neuronal marker protein gene product 9.5 (PGP9.5), and in associated glial cells, detected by glial fibrillary acidic protein (GFAP) and S100 immunostaining (**Fig. 7D**).

PAMAM-Chol-Cy5 is positively charged whereas PEG-Poly-QD is negatively charged. To explore the influence of surface charge on cellular targeting, PAMAM-Chol-Cy5 NPs were synthesized with a negative charge (zeta potential -15±1 mV). Whereas PAMAM-Chol-Cy5(+) was detected mostly in colonocytes and rarely found in neurons (**Fig. S9A, B**), PAMAM-Chol-Cy5(-) was excluded from colonocytes, retained in the lamina propria, and preferentially homed to the nervous system (**Fig. S9C, D**). Thus, a positive surface charge favors epithelial cell delivery whereas a negative surface charge favors mucosal penetration and delivery to the nervous system.

### NP encapsulation enhances antagonism of PAR_2_-induced colonic pain

The availability of NP formulations that would preferentially deliver AZ3451 to colonic epithelial cells and neurons, where PAR_2_ activation induces inflammation and pain (10, 14), allowed examination of the influence of cell and organelle targeting of PAR_2_ antagonists for treatment of inflammation and pain. To determine whether NP-encapsulated PAR_2_ antagonists ameliorate PAR_2_-induced colonic pain, vehicle, PAMAM-Chol-Ø, PAMAM-Chol-AZ, unencapsulated AZ3451 or vehicle was administered into the colonic lumen 30 min before intracolonic administration of 2F (**Fig. 8A**). Abdominal nociception was evaluated by measuring withdrawal responses to stimulation of the abdomen with von Frey filaments (VFF). In vehicle-treated mice, 2F caused a sustained decrease in withdrawal threshold for 6 h, indicating mechanical allodynia (**Fig. 8B-I**). PAMAM-Chol-Ø and PEG-Poly-QD did not affect the response to 2F. PAMAM-Chol-AZ (10 µl, 1, 5, 10 µM AZ3451) caused a dose-dependent and sustained increase in VFF withdrawal threshold, consistent with antinociception (**Fig. 8B-G**). Intracolonic PEG-Poly-AZ (10 µl, 5 µM AZ3451) also inhibited 2F-evoked mechanical allodynia for 6 h (**Fig. 8H, I**). Unencapsulated AZ3451 (10 µl, 1, 5, 10 µM) had a small and transient inhibitory action. Intravenous AZ3451 (5 µM, 200 µl) had no effect on 2F-induced mechanical allodynia (**Fig. S10A**). Thus, NP encapsulation markedly increases the magnitude and duration of analgesia of AZ3451 in the colon.

**Figure 8.**
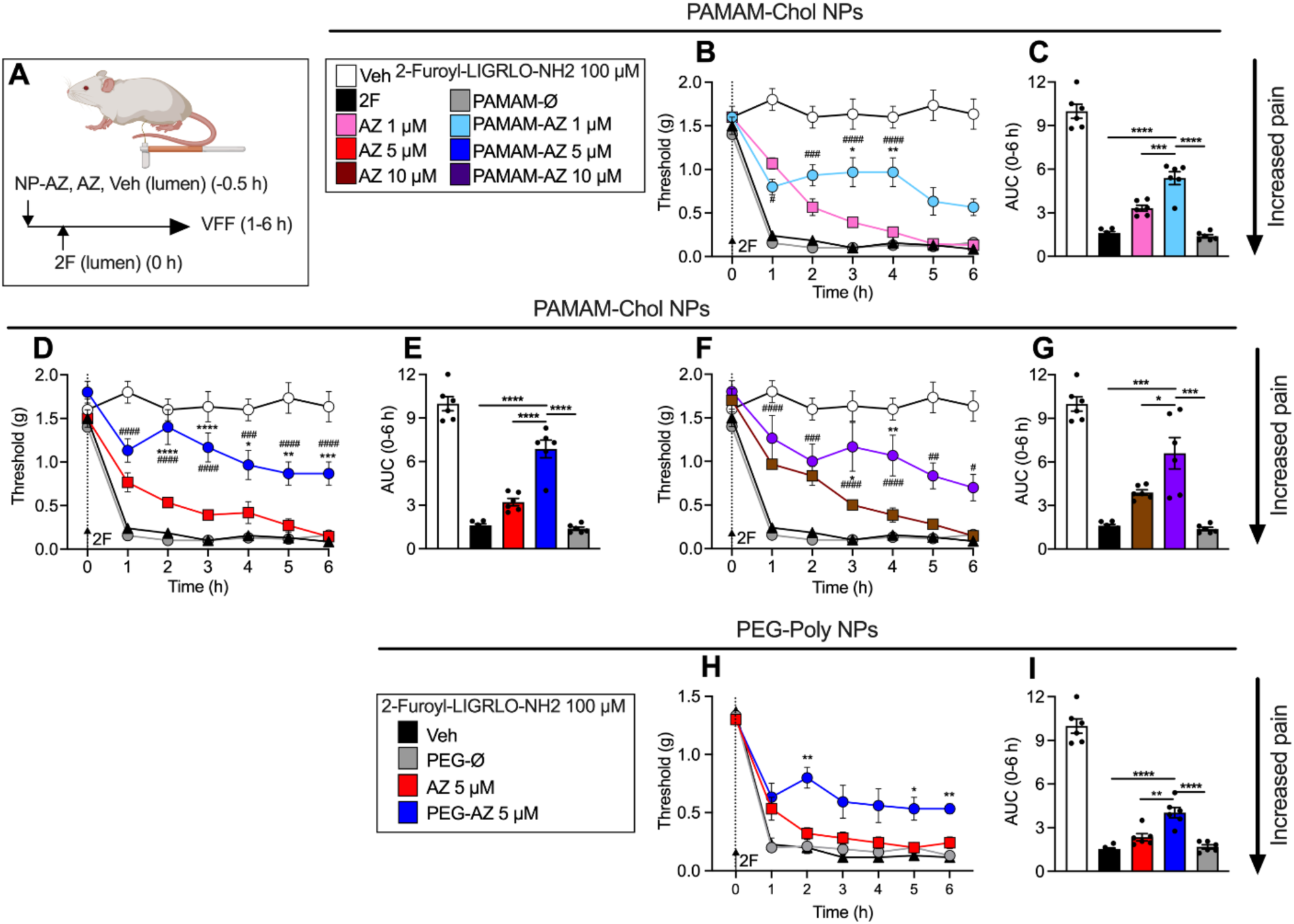
Luminally administered PAMAM-Chol-AZ and PEG-Poly-AZ NPs potently inhibit PAR_2_-induced colonic nociception. **A.** Experimental protocol. **B**-**I.** Abdominal withdrawal threshold to VFF stimulation in mice after intracolonic administration of PAMAM-Chol (**B-G**) or PEG-Poly (**H, I**) followed 30 min later by intracolonic administration of 2F. **B, D, F, H.** Time course. **C, E, G, I**. Area under curve (AUC). Mean±SEM, n=6 mice. **B, D, F, H.** \**P*<0.05, \*\**P*<0.01, \*\*\**P*<0.001, \*\*\*\**P*<0.0001; PAMAM-Chol-AZ3451 (1, 5, 10 µM) or PEG-Poly-AZ3451 vs AZ3451 (1, 5, 10 µM); #*P*<0.05, ##*P*<0.01, ###*P*<0.001, ####*P*<0.0001; PAMAM-Chol-AZ3451 (1, 5, 10 µM) or PEG-Poly-AZ (5 µM) vs 2F (100 µM). One-way ANOVA, Tukey’s multiple comparison test. **C, E, G, I.** \**P*<0.05, \*\**P*<0.01, \*\*\**P*<0.001, \*\*\*\**P*<0.0001. Two-way ANOVA, Tukey’s multiple comparison test.

### NP-encapsulated PAR_2_ antagonist reverses inflammation and pain and normalizes aberrant behavior in preclinical models of IBD

The efficacy of NP-encapsulated PAR_2_ antagonists for the treatment of colonic inflammation and pain was determined in two preclinical mouse models of IBD, induced by administration of trinitrobenzene sulphonic acid (TNBS) or dextran sodium sulphate (DSS). PAMAM-Chol-Ø, PAMAM-Chol-AZ, unencapsulated AZ3451 (150 µl, 5 µM AZ3451) or vehicle was administered into the colon lumen 3 d after intracolonic TNBS, which induces inflammation, pain and endocytosis of PAR_2_-muGFP (14) (**Fig 9A**). TNBS induced weight loss, colonic shortening (**Fig. S11A, B**), abdominal mechanical allodynia (**Fig. 9B, C**), and expression of proinflammatory tumor necrosis factor-alpha (TNFα), interleukin-1beta (IL-1β) and chemokine (C-X-C motif) ligand 1 (CXCL1) mRNA (**Fig. 9D-F**). PAMAM-Chol-Ø had no effect on allodynia or inflammation. PAMAM-Chol-AZ reversed TNBS-induced allodynia and TNFα, IL-1β and CXCL1 expression from 1-6 h. Unencapsulated AZ3451 had no effect on allodynia but modestly inhibited inflammation.

**Figure 9.**
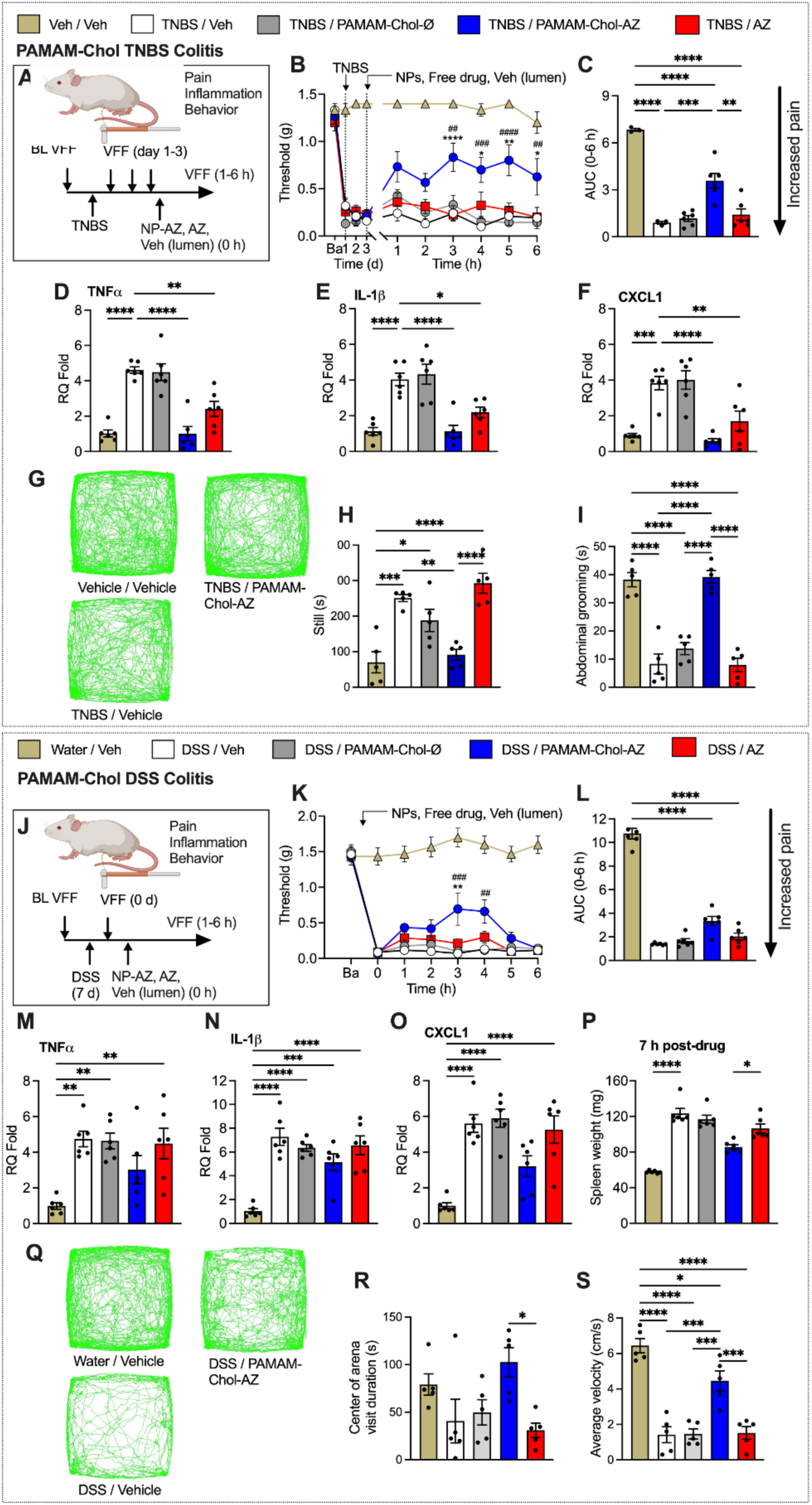
Luminally administered PAMAM-Chol-AZ NPs reverse nociception and inflammation and normalize aberrant behavior in preclinical models of IBD. Nociception and inflammation in mice with TNBS-induced colitis (**A-I**) or DSS-induced colitis (**J-S**)**. A, J.** Experimental protocol. **B, C, K, L.** Abdominal withdrawal threshold to VFF stimulation in mice after intracolonic administration of vehicle, PAMAM-Chol-Ø, PAMAM-Chol-AZ or unencapsulated AZ3451. **B, K.** Time courses. **C, L.** Area under curve (AUC) of time course. **D-F, M-O.** Colonic cytokine and chemokine mRNA levels at 6 h after vehicle, PAMAM-Chol-Ø, PAMAM-Chol-AZ or unencapsulated AZ3451. **P**. Spleen weight at 7 h after vehicle, PAMAM-Chol-Ø, PAMAM-Chol-AZ or unencapsulated AZ3451. **G-I, Q-S**. Spontaneous behavior of mice at 3 h after vehicle, PAMAM-Chol-Ø, PAMAM-Chol-AZ or unencapsulated AZ3451. **G, Q.** Traces of mouse movement over 30 min. **H.** Time spent still. **I.** Time spent grooming abdomen. **R.** Duration of visits to central area. **S.** Average velocity. Mean±SEM, n=5 or 6 mice. **C-F, H-I, L-P, R-S.** \**P*<0.05, \*\**P*<0.01, \*\*\**P*<0.001 \*\*\*\**P*<0.0001. **B,K**. \**P*<0.05, \*\**P*<0.01, \*\*\*\**P*<0.0001, TNBS / PAMAM-Chol-AZ *vs* TNBS / AZ, and ^##^*P*<0.01, ^###^*P*<0.001, ^####^*P*<0.0001 TNBS / PAMAM-Chol-AZ *vs* PAMAM-Chol-Ø. Two-way ANOVA, Tukey’s multiple comparison test. **B, K.** Two-way ANOVA, Tukey’s multiple comparison test. **C-F, H-I, L-P, R-S.** One-way ANOVA, Tukey’s multiple comparison test.

The effects of TNBS colitis on spontaneous behavior were determined using a spectrometer that objectively quantifies behaviors affected by pain (27, 39). TNBS colitis markedly altered locomotor, exploratory and grooming behavior (**Fig. 9G-I, Fig. S12, Movie S1**). TNBS increased the still time and number of still events and decreased abdominal grooming time and events. TNBS decreased average velocity, track length, activity and body length (hunching). Three hours after intracolonic injection, PAMAM-Chol-AZ normalized TNBS-induced changes in still time and events, abdominal grooming time and events, average velocity, track length, activity and body length. PAMAM-Chol-Ø and unencapsulated AZ3451 had no effects.

The efficacy of PAMAM-Chol-AZ was confirmed in a second preclinical IBD model, DSS-induced colitis, which is also associated with PAR_2_ endocytosis (14). DSS caused weight loss, disease symptoms and colonic shortening (**Fig. S11C-E**). PAMAM-Chol-Ø, PAMAM-Chol-AZ, unencapsulated AZ3451 (10 µl, 5 µM AZ3451) or vehicle was administered into the colon lumen at 8 d after beginning DSS treatment (**Fig. 9J**). DSS caused allodynia (**Fig. 9K, L**) and increased TNFα, IL-1β and CXCL1 expression (**Fig. 9M-O**) and spleen weight (**Fig. 9P**). PAMAM-Chol-AZ partially normalized allodynia, cytokine and chemokine expression and spleen weight. PAMAM-Chol-Ø and unencapsulated AZ3451 had no effect.

DSS strikingly affected locomotor, exploratory and grooming behaviors (**Fig. 9Q-S, Fig. S13, Movie S2**). DSS decreased center arena visit times and events, average velocity, track length, activity, distance from the wall, total, body and abdominal grooming events and times, and walk, trot and run events and times. DSS increased still events and time and decreased body length. Within 3 h, PAMAM-Chol-AZ normalized locomotor, exploratory and grooming behavior, and restored body length (reversal of hunching). PAMAM-Chol-Ø and unencapsulated AZ3451 had no effects.

The antinociceptive and anti-inflammatory actions of PEG-Poly-AZ were similarly examined in mice with TNBS- and DSS-evoked colitis. Intracolonic PEG-Poly-AZ (150 µl, 5 µM AZ3451) partially reversed TNBS- and DSS-evoked allodynia (**Fig. 10A-C**, **Fig. 10J-L**) and TNFα, IL-1β and CXCL1 expression (**Fig. 10D-F**, **Fig. 10M-O**). PEG-Poly-AZ restored TNBS- and DSS-induced changes in locomotor, exploratory and grooming behaviors (**Fig. 10G-I, P-R, Fig. S14, Fig. S15, Movie S3, Movie S4**).

**Figure 10.**
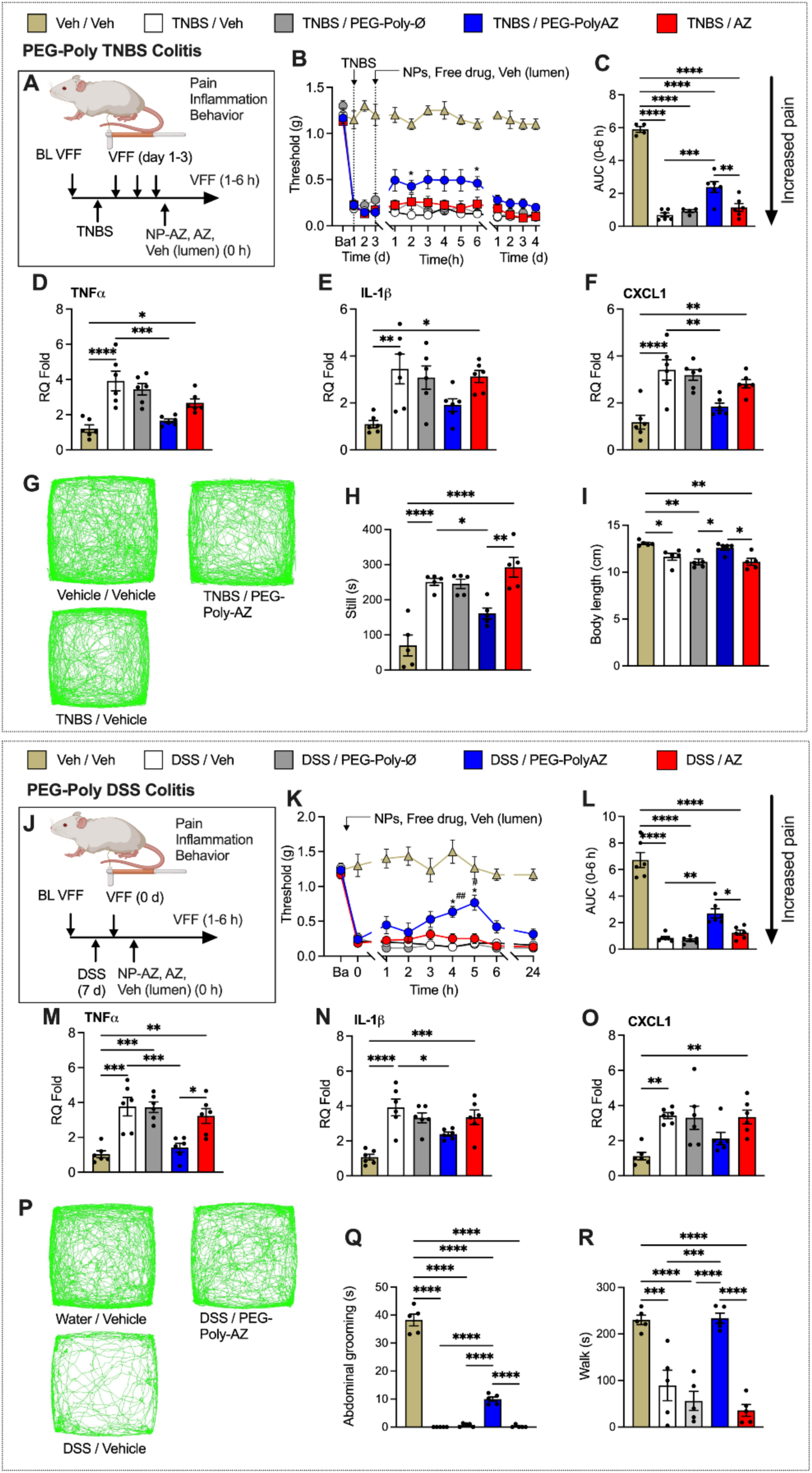
Luminally administered PEG-Poly-AZ NPs reverse nociception and inflammation and normalize aberrant behavior in preclinical models of IBD. Nociception and inflammation in mice with TNBS-induced colitis (**A-I**) or DSS-induced colitis (**J-R**)**. A, J.** Experimental protocol. **B, C, K, L.** Abdominal withdrawal threshold to VFF stimulation in mice after intracolonic administration of vehicle, PEG-Poly-Ø, PEG-Poly-AZ or unencapsulated AZ3451. **B, K.** Time courses. **C, L.** Area under curve (AUC) of time course. **D-F, M-O.** Colonic cytokine and chemokine mRNA levels at 6 h after vehicle, PEG-Poly-Ø, PEG-Poly-AZ or unencapsulated AZ3451. **G-I, P-R**. Spontaneous behavior of mice at 3 h after vehicle, PEG-Poly-Ø, PEG-Poly-AZ or unencapsulated AZ3451. **G, P.** Traces of mouse movement over 30 min. **H.** Time spent still. **I.** Body length. **Q.** Time spent grooming abdomen. **R.** Time spent walking. Mean±SEM, n=5 or 6 mice. **C-F, H-I, L-O, Q-R**. \**P*<0.05, \*\**P*<0.01, \*\*\**P*<0.001 \*\*\*\**P*<0.0001. **B,K**. \**P*<0.05 TNBS / PEG-Poly-AZ vs TNBS / AZ, and ^#^*P*<0.05, ^##^*P*<0.01 TNBS / PEG-Poly-AZ vs TNBS / PEG-Poly-Ø. Two-way ANOVA, Tukey’s multiple comparison test. **B, K.** Two-way ANOVA, Tukey’s multiple comparison test. **C-F, H-I, L-O, Q-R.** One-way ANOVA, Tukey’s multiple comparison test.

Intravenous AZ3451 (5 µM, 200 µl) had no effect on TNBS-induced mechanical allodynia (**Fig. S10B**).

Thus, locally administered NPs that deliver PAR_2_ antagonists to endosomes of colonic epithelial cells and the nervous system normalize inflammation, nociception and aberrant behaviors in two preclinical models of IBD in which mammalian and bacterial proteases drive disease. Unencapsulated antagonist, whether administered locally or systemically, is ineffective.

## Discussion

In this study we have shown that NPs deliver a PAR_2_ antagonist to endosomes of epithelial cells and neurons of the colon resulting in a transformational increase in efficacy for the treatment of pain, inflammation and associated aberrant behaviors in preclinical mouse models of IBD. Enhanced therapeutic activity is attributable to effective antagonism of preexisting PAR_2_ signals in organelles. A non-encapsulated antagonist of PAR_2_ is minimally efficacious in IBD models because it is unable effectively reverse ongoing PAR_2_ signals in organelles. We have thus identified an effective and long-lasting therapeutic approach based on the local administration of nanomedicines designed to deliver antagonists to intracellular sites of signaling in the key cell types that control pain in the colon. The approach may enhance the efficacy of GPCR-targeted therapeutics for diverse diseases.

### Antagonism of PAR_2_ signaling in organelles provides efficacious and long-lasting relief from pain

Because GPCRs are considered cell surface receptors, antagonists are generally evaluated for their ability to prevent GPCR activation at the plasma membrane. Whether antagonists terminate preexisting GPCR signals in organelles, which requires penetration of plasma and organelle membranes and engagement of GPCRs within multi-protein signaling complexes of acidified organelles, is seldom tested. Our results show that a PAR_2_ antagonist encapsulated into dendrimer or polymeric NPs rapidly and completely terminates the activation of PAR_2_, mGαq and βARR1 in early, late and recycling endosomes and the *cis*- and *trans*-Golgi apparatus, and fully reverses the activation of ERK, a key PAR_2_ effector, in the cytosol and nucleus. In sharp contrast, the unencapsulated antagonist has minor and delayed effects or is inactive, which suggests that it is unable to penetrate the plasma and organelle membranes to antagonize preactivated PAR_2_. The accumulation of NPs in early and late endosomes explains the effective antagonism of PAR_2_ signaling in these compartments. Once released from NPs in early and late endosomes, the antagonist diffuses throughout the cell and thereby antagonize PAR_2_ in recycling endosomes, the Golgi apparatus and at the plasma membrane, which we propose contributes to the prolonged analgesia.

Our results show that NPs containing a PAR_2_ antagonist, when administered locally into the colon, effectively relieve pain and inflammation and restore aberrant behavior in mouse models of IBD. NPs that deliver the antagonist to colonocytes (dendrimers) or colonic neurons (core-shell polymers), prominent sites of PAR_2_ expression (14), are both efficacious. In contrast, unencapsulated PAR_2_ antagonist, whether administered locally or systemically, is largely inactive. Delivery of the NP-encapsulated antagonist to sites of PAR_2_ pain signaling in organelles of relevant cell types accounts for the transformative therapeutic benefits of NPs developed in the current study. We propose that NP-encapsulated PAR_2_ antagonists are a much-needed non-opioid therapy for abdominal pain that is a hallmark of digestive diseases (40). Coupling these nanomedicines with an oral delivery strategy would expand clinical utility. Although NPs were not toxic to cultured cells, *in vivo* toxicology and pharmacokinetic analyses are required to advance NP-encapsulated PAR_2_ antagonists to clinical trials of digestive diseases. Further studies are required to understand the basis of NP tissue penetration, cell-specific targeting, and uptake and retention in endosomes, which could further improve the efficacy and duration of the protective effects, since AZ3451 is released from NPs over several days *in vitro*.

Antagonism of PAR_2_ signals in organelles is necessary for effective therapy because PAR_2_ redistributes from the plasma membrane to endosomes of colonocytes and nociceptors in the inflamed colon (10, 14), which is attributable to protease activation (38). Furthermore, PAR_2_ signaling in endosomes of colonocytes and nociceptors mediates inflammation and pain (10, 14, 17). We leveraged the propensity of NPs to accumulate in endosomes to deliver an antagonist to target disease-relevant PAR_2_ signaling in organelles. AZ3451 release from dendrimer and core-shell NPs is not pH-dependent, suggesting that pH-dependent cargo release in acidified endosomes is unlikely to contribute to persistent efficacy, in contrast to pH-tunable soft polymer NPs (26–28). pH-responsive NPs are not suitable for the treatment of inflammatory pain because they would be expected to prematurely disassemble before cellular uptake due to the acidity of extracellular fluid in inflamed tissues, including the colon (29).

Our results show that antagonism of PAR_2_ signals in relevant cells is also necessary for effective treatment. Dendrimer NPs accumulate in colonocytes whereas core-shell NPs penetrate the mucosa and accumulate in neurons and glial cells. Antagonist delivery to endosomes of colonocytes would prevent PAR_2_-induced paracellular permeability and inflammation and delivery to neurons would block sensitization and pain. The mucosal retention of dendrimers is likely due to their soft structure and cationic surface charge, rendering them mucoadhesive (33, 41). Accumulation of dendrimers in endosomes of colonocytes is attributable to the positive surface charge that enhances interactions with negative cell surfaces (41, 42). In contrast, the slightly negative core-shell NPs penetrate deeply into the mucosa, submucosa and musculature, gaining access to neurons. Mucus and tissue penetration is consistent with previous studies of PEGylated NPs (43). While colonocyte and neuronal endosomal uptake is important for efficacy, further development of NPs by surface conjugation of targeting moieties to enhance delivery to specific cell types, and core optimization to maximize drug loading and sustained endosomal release are required to advance such formulations to the clinic.

A striking finding of the current study is that NP-encapsulated PAR_2_ antagonist restores pathologic locomotor, exploratory and grooming behaviors in mice with IBD, whereas unencapsulated antagonist is ineffective. Although TNBS and DSS colitis affected behavior to different extents, dendrimer and core-shell NPs containing a PAR_2_ antagonist were effective in both models. The restoration of behavior is likely attributable to the anti-inflammatory and analgesic properties of the NP-encapsulated antagonist in the colon. Abdominal pain was evaluated in the current study by measuring withdrawal responses of the mice to abdominal stimulation with calibrated filaments, with a decreased withdrawal threshold reflecting abdominal mechanical allodynia. This approach avoids the confound of surgical placement of electrodes to measure visceromotor responses, but does not assess responses to direct stimulation of the colon. Although abdominal mechanical allodynia and reduced locomotion, exploration and abdominal grooming are all consistent with colonic inflammation and pain, future studies that specifically examine mechanosensitivity of the colon are required.

### Implications for local delivery of antagonists for reversal of intracellular signaling

Our finding that a NP-encapsulated PAR_2_ antagonist terminates signaling in organelles and provides efficacious and long-lasting analgesia has broad implications. GPCRs are the single largest target of approved drugs (2). Although modification of the lipophilicity and acidity of small molecule GPCR antagonists can enhance antagonism of intracellular signals and improve therapeutic efficacy(24), the criteria for development of endosomally-targeted antagonists are not fully defined. Encapsulation into NPs would facilitate antagonist delivery to sites of disease-relevant GPCR signals in organelles and thereby improve therapeutic efficacy. The local administration of NPs for targeting receptors in endosomes confers further advantages. For applications that require therapeutics to reach cytoplasmic, nuclear or mitochondrial targets, the necessity and obstacles of endosomal escape restrict the usefulness of NP-mediated organelle targeting strategies (44). The use of NPs to deliver therapeutics to endosomal targets overcomes this limitation. NPs engineered to target and persist in diseased tissues, cells and organelles and to exhibit sustained release of drug cargo offer the prospect of improved therapeutic efficacy whilst minimizing dosing and side effects from systemic exposure (45). However, systemically administered NPs face significant delivery hurdles including opsonization, phagocytosis and sequestration in the liver and spleen, NP surface fouling *via* a protein corona that impedes tissue targeting, and the challenge of selectively targeting NPs to organelles of diseased cells. The local administration of NPs encapsulating antagonists of endosomally-enriched GPCR targets surmounts the obstacles of systemic delivery and is applicable to treatment of many diseases.

## Materials and Methods

Complete Materials and Methods are provided in Supporting Information.

### Mice

Institutional Animal Care and Use Committees approved all studies. Male C57BL/6J mice (8-12 weeks) were from JAX® or Queens University. *Par_2_-mugfp* knockin mice have been described (14).

### Synthesis of PAMAM-Chol NPs

Positively charged PAMAM-Chol(5) was prepared as described (30). Supporting Information describes encapsulation of AZ3451 into PAMAM-Chol(5) NPs, Cy5 coupling to PAMAM-Chol, and preparation of negatively charged PAMAM-Chol(5).

### Synthesis of PEG-Poly NPs

PEG-Poly-AZ NPs were synthesized by flash nanoprecipitation (FNP) as described (46). Supporting Information describes encapsulation of QDs into core-shell NPs *via* FNP.

### NP characterization

NPs were characterized by measurement of hydrodynamic diameter and zeta potential, by transmission electron microscopy (TEM), and by determination of total NP solids concentration (47).

### AZ3451 loading and release

AZ3451 loading into NPs and the time course of release were measured by reverse-phase HPLC.

### Uptake and trafficking of fluorescent NPs

#### HEK293T cells, T84 cells, DRG neurons

HEK293T cells were transduced with CellLight Rab5a-RFP or Rab7a-RFP to identify early and late endosomes, respectively. HEK293T cells, T84 cells or DRG neurons were incubated with PAMAM-Chol-Cy5 or PEG-Poly-QD for indicated times at 37°C. HEK293T cells were directly imaged. T84 cells and DRG neurons were fixed (4% paraformaldehyde, 20 min, on ice) and processed to detect EEA1 or NeuN by immunofluorescence.

#### Colon

PAMAM-Chol-Cy5 or PEG-Poly-QD was administered into the colon of sedated mice, including *Par_2_-mugfp* mice) (14), by enema. The colon was collected after various times, fixed (4% paraformaldehyde, overnight, 4°C), and frozen sections were prepared. The neuronal marker PGP9.5 and the glial cell markers GFAP and S100 were detected by indirect immunofluorescence.

#### Confocal microscopy

Specimens were imaged using a Leica SP8 confocal microscope with 40x or 63x objective (NA = 1.4).

### cDNAs

Supporting Information described cloning and sources of cDNAs for BRET assays.

### BRET assays

#### NbBRET

NbBRET biosensors were expressed in HEK293T cells in which endogenous PAR_2_ was deleted (48). Supporting Information describes transfection of nbBRET sensors into HEK293T and T84 cells, the treatment of cells with PAR_2_ agonists and antagonists, and the nbBRET assay protocol. To determine whether NPs can reverse PAR_2_ activation, cells were challenged with 2 F or trypsin. Seven min after agonist challenge, cells were incubated with vehicle, PAMAM-Chol-AZ, PEG-Poly-AZ or unencapsulated AZ3451 and BRET was recorded for an additional 30 min.

#### EbBRET

Supporting Information describes transfection of ebBRET sensors into HEK293T cells, the treatment of cells with PAR_2_ agonists, and the ebBRET assay protocol (20, 21).

#### GEMTA

Supporting Information describes transfection of GEMTA sensors into HEK293SL cells for measurement of Gαq activation in subcellular compartments (25). To determine whether NPs can reverse Gαq activation, cells were challenged with 2F or trypsin. Two min after agonist challenged, cells were exposed to vehicle, PAMAM-Chol-AZ, PEG-Poly-AZ or unencapsulated AZ3451 and BRET was recorded for an additional 20 min.

#### BRET measurements

Supporting Information describes BRET measurements.

### FRET assays

Supporting Information describes transfection of HEK-PAR_2_KO cells with FLAG-PAR_2_-HA and nucEKAR or cytoEKAR FRET sensors and FRET measurements (8). Seven min after agonist stimulation, cells were incubated with vehicle, PAMAM-Chol-AZ, PEG-Poly-AZ or unencapsulated AZ3451 and FRET was recorded for an additional 30 min.

### Patch clamp recording

Supporting Information describes dispersion and culture of mouse DRG neurons, patch clamp measurements of rheobase (10), and treatment of neurons with PAR_2_ agonists and NPs.

### Preclinical colitis models

Supporting Information describes the intracolonic administration of PAR_2_ agonist 2F. Colitis was induced by intracolonic administration of 2,4,6-trinitrobenzene sulphonic acid (TNBS) (14) or by administration of dextran sodium sulphate (DSS) in drinking water for 7 days (14). PAMAM-Chol-Ø, PAMAM-Chol-AZ3451, PEG-Poly-Ø, PEG-Poly-AZ, unencapsulated AZ3451 or vehicle was injected into the colon by enema 30 min before 2F, or 96 h post TNBS, or 8 d post commencing DSS treatment. AZ3451 was also administered intravenously.

### Colonic nociception

Withdrawal responses to stimulation of the abdomen with von Frey filaments (VFF) were recorded to assess visceral pain-like behavior as described (14). Investigators were blinded to treatments.

### Spontaneous non-evoked behavior

Spontaneous behavior was assessed using a spectrometer that objectively quantifies locomotor, exploratory and grooming behavior (27, 39).

### Colonic inflammation

Cytokine and chemokine expression in the colon was measured by qRT-PCR as described (14).

### Statistical analysis

Results were analyzed and graphs prepared using Prism 10. Data sets were normality tested with a D’Agostino & Pearson and Shapiro-Wilk test and analyzed statistical significance using one- or two-way ANOVA or Kruskal-Wallis test with Tuckey’s, Šidák’s, Bonferroni’s or Dunn’s post hoc test. Results are expressed as mean±SEM. The level of significance was set at up *P* ≤ 0.05.

## Acknowledgments

N.A. Lambert (Augusta University) kindly provided cDNA encoding mGα isoforms coupled to Venus. A. Thomsen (New York University) kindly provided cDNA encoding CAAX and FYVE. We thank Ana Manu for technical assistance. This research was funded by National Institutes of Health Grants R01NS102722 (NWB, BLS), R01DK118971 (NWB, BLS), R01DE026806 (NWB, BLS), R01DE029951(NWB, BLS, KL), RM1DE033491 (NWB, BLS, KL), R01DK130293 (MB, NWB), and 5R01NS125413 (DDJ), and by Department of Defense Grants W81XWH1810431 (NWB, BLS) and W81XWH2210239 (NWB, BLS)

## Materials and Methods

### Materials

Tetrahydrofuran (THF, high pressure liquid chromatography, HPLC, grade), acetonitrile (ACN, HPLC grade), and Tween 80 (cell culture grade) were from Fisher Scientific (USA). Dimethyl sulfoxide (DMSO, HPLC grade), polylactic acid homopolymer (PLA, 10-18 kD), rubrene (98%), PAMAM-G3, cholesterol chloroformate, N,N-diisopropylethylamine, dichloromethane, BODYPY (BDP) 650/655 and cyanine 5 (Cy5) were from Sigma-Aldrich (USA). Poly(lactic acid)-poly(ethylene glycol) (PEG-PLA, 5k-5k) was donated by Evonik Inc (USA). Quantum dots (660 nm emission) were from Ocean Nanotech. AZ3451 (99.6%) was from MedChemExpress (USA). Ultra-pure water (18.2 MΩ·cm) was generated by a MilliporeSigma Milli-Q Water Purification System (USA).

### Mice

Institutional Animal Care and Use Committees of New York University and Queens University approved all studies. Male C57BL/6J mice (8-12 weeks) were from JAX® (#000664 JAX®, wild type) or Queens University. Knockin mice expressing PAR_2_ C-terminally fused to monomeric ultrastable green fluorescent protein (PAR_2_-muGFP) have been described (1). Mice were maintained in a light (12 h light/dark cycle) and temperature (22°C) controlled environment with *at libitum* access to food and water. Mice were randomly assigned to experimental groups using Research Randomizer (www.randomizer.org). Randomization avoided operator bias and ensured that experimental groups were composed of animals from different cages to avoid cage-related issues. Investigators were blinded to treatment for studies in mice. Test agents were prepared by a technician and placed into coded tubes. For the behavioral assays, mice were positioned in the apparatus by a technician and the investigator was blinded to the treatment.

## Methods

### Synthesis of positively charged PAMAM-Chol NPs

#### Polymer synthesis

PAMAM-Chol(5) was prepared by incubating PAMAM-G3 (15 µmol) with cholesterol chloroformate (75 µmol) and N,N-diisopropylethylamine (300 μmol) in 5 mL dichloromethane under stirring at room temperature for 24 h, followed by dialysis in ultrapure water for 72 h (2).

#### PAMAM-Chol-AZ

PAMAM-Chol(5) (1 mg) and AZ3451 (200 µg) were dissolved in 100 µL chloroform; 2 mL water was added and the mixture was sonicated for 2 min. An additional 5 mL of water was added and excess solvent was removed by rotary evaporation. Unencapsulated AZ3451 was eliminated by centrifugation (3000 rpm, 30 min). NPs were lyophilized with 5% trehalose.

#### Fluorescent PAMAM-Chol NPs

Cy5 conjugated to N-hydroxysuccinimide (*NHS*) were coupled to surface amine groups of PAMAM-Chol. PAMAM-Chol NPs were mixed with BDP-NHS ester or Cy5-NHS ester (Lumiprobe) in a 1:50 NP:dye ratio and mixed overnight at room temperature. The suspension was placed in a 3500 MWCO dialysis bag and dialyzed against water to remove excess dye.

### Synthesis of negatively charged PAMAM-Chol NPs

#### Polymer synthesis

To obtain negatively charged PAMAM-Chol(5), PAMAM-G2.5 (63 mg, 10 μmol) was dissolved in 5 mL of DMSO. EDC (48 mg, 250 μmol) and NHS (12 mg, 100 μmol) were added to the solution, which was stirred for 30 min. Then, 1-dodecylamine (9.3 mg, 50 μmol) was added, and the mixture was stirred at room temperature overnight and then dialyzed against water for 72 h (2).

#### Fluorescent PAMAM-Chol NPs

Cy5 conjugated to an amine group was coupled to surface carboxylate groups of negative PAMAM-Chol. PAMAM-Chol NPs were mixed with NH_2_-Cy5 (Lumiprobe) in a 1:50 NP:dye ratio and mixed overnight at room temperature. The suspension was placed in a 3500 MWCO dialysis bag and dialyzed against water to remove excess dye.

### Synthesis of PEG-Poly NPs

#### PEG-Poly-AZ assembly

PEG-Poly-AZ NPs were synthesized by flash nanoprecipitation (FNP) (3). An organic stream of 75 vol % THF and 25 vol % DMSO containing dissolved AZ3451 (3.2 mg/mL), polylactic acid (PLA, 16.8 mg/mL) and PEG-PLA (20 mg/mL) was rapidly mixed using a multi-inlet vortex mixer against 3 ultra-pure water streams of equal volume. The mixture was collected in a quenching water bath to reduce the final organic concentration to 5 vol %. To remove organic solvent and unencapsulated drug, the NP solution was dialyzed at 4°C against a 100-fold excess volume of ultra-pure water using a 12-14 kD molecular weight cut-off (MWCO) regenerated cellulose dialysis membrane (Spectra/Por, Spectrum Laboratories); water was changed hourly for 6 h. For cell-based and animal studies, NP solutions were sterile filtered using a 0.22 µm nylon syringe filter (Fisher Scientific).

#### Fluorescent PEG-Poly NPs

Quantum dots (QDs) were encapsulated into core-shell NPs *via* FNP. Organic streams containing dissolved PEG-PLA (6 mg/mL), PLA (13 mg/mL) and QDs (1 mg/mL) were prepared in THF and rapidly mixed against equal volumes of ultrapure water in a confined impinging jet mixer. The mixture was collected in an aqueous quench bath to reduce the final organic solvent concentration to 10 vol %. The NP solution was dialyzed against ultrapure water as described above. For cell-based and animal studies, NPs were assembled using sterile materials and following aseptic techniques.

### NP characterization

The hydrodynamic diameter and zeta potential of the NPs were measured with a Malvern Nano ZS90 Zetasizer. NPs were observed by transmission electron microscopy (TEM) using a FEI Talos L120C microscope. To prepare TEM samples, 0.5-1.0 mg/mL NP solutions were drop-cast onto ultrathin carbon on lacey carbon, 300 mesh Au TEM grids (TedPella) and allowed to dry. Samples were negatively stained with 2% phosphotungstic acid (Sigma Aldrich). Total NP solids concentration was determined using a thermogravimetric analyzer (TGA, Discovery 550 TGA, TA Instruments). NPs in solution were heated under nitrogen from 25°C to 105°C at a rate of 10°C/min and held at 105°C for 20 minutes to ensure complete water evaporation. Total NP concentration was determined by dividing the final solid weight by the starting sample volume (4).

### AZ3451 loading

AZ3451 loading into NPs was measured by reverse-phase HPLC using either an Agilent 1260 Infinity HPLC and an HC-C18(2) column (4.6 x 150 mm, 4 µm pore size) or a ThermoFisher Vanquish Core HPLC and a Hypersil Gold C18 column (4.6 mm x 150 mm, 75 Å pore size). The mobile phase was water and acetonitrile with 0.1% trifluoracetic acid and a flow rate on 0.5 or 1 mL/min. Absorbance at 254 or 319 nm was measured. AZ3451 was quantified by comparing absorbance to a standard curve.

### AZ3451 release

AZ3451 release was measured by HPLC. NPs were dispersed in PBS to a solids concentration of 1.1 mg/mL. The suspension (1 mL) was placed in a cellulose dialysis bag (Spectra/Por 3, MWCO 3500, Spectrum) and dialyzed against 10 mL of release medium (a mixture of 0.1 M citric acid and 0.2 M Na_2_HPO_4_ buffer solutions containing 0.1% w/v Tween 80, pH = 7.4, 6.5, 5.5) with shaking at 37°C. At specified times, 0.4 or 1 mL of release media was sampled and the dialysis bag was transferred to a new tube of release medium. Samples were deformulated by adding 100 µL acetonitrile and analyzed for AZ3451 by HPLC.

### NP toxicity assay

Cytotoxicity of NPs in HEK293T cells was evaluated using a CCK-8 kit (Thermo Fisher Scientific, Cat# NC9624837). Cells were plated on clear 96-well plates. After 24 h, cells were incubated with vehicle (negative control), 0.2% Triton-X-100 (positive control), PAMAM-Chol-Ø (0.3-10 µg/mL) or PEG-Poly-Ø (0.3-10 µg/mL) for 4 or 24 h. A 1:10 dilution of CCK-8 reagent: DMEM was added to each well and incubated for 30 min h at 37°C. The absorbance of the plate was measured at 450 nm with a CLARIOstar Microplate Reader (BMG Labtech). Absorbance readings were normalized to that of the vehicle-treated cells.

### Uptake and trafficking of fluorescent NPs

#### HEK293T cells

HEK293T cells were plated on 35 mm glass bottom dishes (MatTek) pre-coated with poly-D-lysine and incubated in DMEM/10% FBS. After 24 h, cells were transduced with BacMam 2.0 CellLight Early Endosomes-RFP (4 μL/dish) and BacMam 2.0 CellLight Late Endosomes-GFP (4 μL/dish) (ThermoFisher). Cells were incubated overnight then washed twice with FluoroBrite^TM^ DMEM. Cells were incubated with PAMAM-Chol-Cy5 (3 µg/mL) or PEG-Poly-QD (13.5 µg/mL) NPs. Live cells were imaged live at 0 h, 2 h and 4 h using the Leica SP8 confocal microscope with LAS X imaging software. NP uptake to early and late endosomes was quantified using ImageJ (NIH). Regions of interest (ROIs) were generated using the early or late endosomal channels (GFP or RFP). A threshold of the endosome channel was set using the minimum algorithm with a range of 25000-65535 relative fluorescence units (RFUs). ROIs were then overlayed to the NP channel and the mean RFU was measured for each ROI. NP-positive endosomes were defined as an endosome (ROI) with a mean RFU >1200 (maximal RFU recorded at the 0 h time point) in the NP channel and expressed as a percentage of the total number of endosomes in each cell per dish. <u>T84 cells and DRG neurons.</u> T84 cells or DRG neurons in short-term culture were incubated in FluoroBrite^TM^ DMEM containing PAMAM-Chol-Cy5 or PEG-Poly-QD (1.5 mg/mL) for 30 min, 1, 2, 4 and 8 h at 37°C. T84 cells and DRG neurons were fixed with 4% paraformaldehyde in PBS on (20 min, on ice). Cells were washed, incubated in blocking buffer (PBS, 5% normal horse serum, 0.03% Saponin) (1 h, room temperature). Cells were incubated with the primary antibodies: mouse anti-EEA1 (1:100, early endosomal marker, 1G11, Ab70521, AbCam) or mouse anti-NeuN (1:1000, neuronal marker, ABN90, Millipore, Burlington MA) (overnight, 4° C). Cells were washed and incubated with the secondary antibodies: donkey anti-mouse Alex647 (1:1000, ThermoFisher, Waltham MA) (room temperature, 1 h). Cells were washed and mounted using ProLong^TM^Glass Antifade (ThermoFisher) hard set mounting media.

#### Colon

PAMAM-Chol-Cy5 or PEG-Poly-QD (150 µl, 1 mg/mL) was administered into the colon of sedated (3% isoflurane) mice by enema (3 cm from anus). The colon was collected after 1, 2 or 8 h, and immersion fixed in 4% paraformaldehyde (overnight, 4°C). Tissues were washed, cryoprotected in sucrose (30%, 48 h, 4°C), embedded in tissue freezing medium (TFM, General Data), and frozen sections were prepared. Sections were incubated with DAPI to stain nuclei. To assess the uptake of PAMAM-Chol-Cy5 NPs into colonocyte endosomes containing PAR_2_, PAMAM-Chol-Cy5 was administered into the colon of knockin mice expressing PAR_2_ C-terminally fused to monomeric ultrastable green fluorescent protein (*Par_2_-mugfp* mice) (1). PAR_2_-GFP and EAA1 were detected as described (1). To assess the uptake of PEG-Poly-QD NPs into neurons and glial cells, colonic sections from mice treated with PEG-Poly-QD were incubated with rabbit anti-PGP9.5 (neuronal marker, Abcam, Ab109261), rabbit anti-GFAP (glial marker, Abcam, Ab290) or S100 (glial marker Abcam Ab52642) (overnight at 4°C). Sections were washed and incubated with donkey anti-mouse Alexa 647 (ThermoFisher) (45 min, room temperature). Sections were mounted with Prolong-Glass (ThermoFisher). To quantify NP retention in the mucosa or muscle layers (including enteric neurons) of the colon, tissue regions were defined and areas were recorded. The Otsu threshold function of ImageJ was used to identify positive ROIs for NPs and the average relative fluorescence was quantified for each ROI. The mean RFUs for each section was normalized to the area of the tissue section. Data are presented as the mean RFU of all ROIs in a section divided by the area of that section (mean RFU/µm^2^).

#### Confocal microscopy

Specimens were imaged using a Leica SP8 confocal microscope with 40x or 63x objective (NA = 1.4). Images were processed with ImageJ (NIH).

#### cDNAs

mGα coupled to Venus were provided from N.A. Lambert (Augusta University). For nbBRET studies, PAR_2_-NatP was cloned using the Gibson Assembly with cDNA encoding PAR_2_ in a pcDNA5/FRT vector. The 13 amino acid natural peptide fragment of nanoluciferase (GVTGWRLCERILA, NatP) was added to the C-terminus of PAR_2_ with a flexible linker (LRPLGSSGGG). Localization markers CAAX, FYVE, Rab4a, Rab7a, Rab11a, Giantin and TGN38 were tagged on the N-terminus with HA, a short linker (GGSG) and the LgBiT tag while TGN38 was tagged on the C-terminus with HA, a short linker (CGSG) and the LgBiT tag. CAAX and FYVE were from A. Thomsen (New York University).

### BRET assays

#### NbBRET

NbBRET biosensors were expressed in HEK293T cells in which endogenous PAR_2_ was deleted by CRISPR/Cas9 gene editing (HEK-PAR_2_-KO cells) (5). For nbBRET studies of PAR_2_ activation, HEK-PAR_2_KO cells in a 96-well plate were transfected using PEI (Polysciences; 1:6 DNA:PEI) with human PAR_2_-NatP (10 ng/well), HA-LgBiT-CAAX, HA-LgBiT-FYVE, HA-LgBiT-Rab7a, HA-LgBiT-Rab4a, HA-LgBiT-Rab11a, HA-LgBiT-Giantin or TGN38-HA-LgBiT (15 ng/well), and Venus-mGαq, Venus-mGαi, Venus-mGαs or YFP-βARR1 (10 ng/well). After 48 h, cells were washed with HBSS containing 10 mM HEPES at pH 7.4. T84 cells were transfected using Lipofectamine 3000 with POMC-Flag-PAR_2_-NatP (100 ng/well, 96-well plate), HA-LgBiT-CAAX, HA-LgBiT-FYVE, HA-LgBiT-Rab7a, HA-LgBiT-Rab4a, HA-LgBiT-Rab11a, HA-LgBiT-Giantin or TGN38-HA-LgBiT (150 ng/well), and Venus-mGαq (150 ng/well). In some nbBRET experiments, cells were pre-incubated in HBSS (30 min, 37°C) containing 0.45 M sucrose, 1 µM AZ3451 or vehicle (control). To determine whether pretreatment of cells with NPs can prevent PAR_2_ activation, cells were preincubated with vehicle, PAMAM-Chol-Ø, PAMAM-Chol-AZ, PEG-Poly-Ø, PEG-Poly-AZ or unencapsulated AZ3451 (100 pM-3 µM AZ, PAMAM-Chol 0.0003-9.09 µg/mL; PEG-Poly, 0.00078-23.5 µg/mL) for 4 h at 37°C. The substrate for nbBRET (furimazine, 10 µM, 15 min) was added to cells. Baseline was measured for 3 min and cells were then challenged with 2F (10 µM), trypsin (100 nM) or vehicle. BRET was measured for an additional 20 min. To determine whether NPs can reverse PAR_2_ activation, cells were challenged with 2 F (10 µM) or trypsin (100 nM). Seven min after agonist challenge, cells were incubated with vehicle, PAMAM-Chol-AZ, PEG-Poly-AZ or unencapsulated AZ3451 (1 or 3 µM AZ) and BRET was recorded for an additional 30 min.

#### EbBRET

For measurement of Gα and βARR recruitment to subcellular compartments, HEK293T cells in a 96-well plate were transfected with mGαi or mGαq fused to RLuc8, or βARR1 fused to RlucII (BRET donors) (5 ng/well) and FLAG-PAR_2_-HA (10 ng/well) and plasma marker CAAX fused to RGFP (BRET acceptor) or compartments markers (Rab5a, Rab7a, Rab4a, Rab11a, Giantin) fused to tdRGFP (20 ng/well) using PEI (1:4 DNA:PEI) (6, 7). After 48 h, cells were incubated in HBSS-HEPES. Prolume purple (2 μM, Nanolight Technology) was added 6 min before baseline measurements and stimulation with 2F (10 µM) or trypsin (100 nM).

#### GEMTA

For measurement of Gαq activation in subcellular compartments, HEK293SL cells in a 96-well plate were transfected with p63-RhoGEF fused to RlucII (1 ng/well) and FLAG-PAR_2_-HA (10 ng/well) and plasma marker CAAX fused to RGFP (BRET acceptor) or compartments markers (Rab5a, Rab7a, Rab4a, Rab11a, Giantin) fused to tdRGFP (20 ng/well) using PEI (8). After 48 h, cells were incubated in HBSS-HEPES. Prolume purple (2 μM) was added 6 min before baseline measurement. To determine whether NPs can reverse Gαq activation, cells were challenged with 2F (10 µM) or trypsin (100 nM). Two min after agonist challenged, cells were exposed to vehicle, PAMAM-Chol-AZ, PEG-Poly-AZ or unencapsulated AZ3451 (1 or 3 µM AZ) and BRET was recorded for an additional 20 min.

#### BRET measurements

BRET and luminescence were recorded in a Synergy Neo2 Microplate reader (BioTek) using BRET1 filters for nbBRET (donor filter: 460 nm, acceptor filter: 540 nm) and BRET2 filters for ebBRET and GEMTA (donor filter: 410nm, acceptor filter: 515nm). ΔBRET represents the BRET signal in the presence of agonist, minus the BRET signal over time in the presence of vehicle. Data are shown as mean±SEM. from independent experiments as the mean value from triplicate wells.

#### FRET assays

To examine whether NPs can reverse PAR_2_-induced ERK activation, HEK-PAR_2_KO cells in a poly-D-Lysine-coated 96-well plate were transfected with FLAG-PAR_2_-HA (55 ng/well) and nucEKAR or cytoEKAR (40 ng/well) using PEI (1:6 DNA:PEI) (9). Cells were serum restricted overnight before FRET measurement. After 48 h, cells were incubated in HBSS-HEPES. FRET was measured in a CLARIOstar Plus plate reader (BMG Labtech) using cyan (470 nM) and yellow fluorescent protein (535 nM) emission ratios. Baseline was measured for 5 min and then cells were challenged with 2F (10 µM) or vehicle. Seven min after agonist stimulation, cells were incubated with vehicle, PAMAM-Chol-AZ, PEG-Poly-AZ or unencapsulated AZ3451 (1 or 3 µM AZ) and FRET was recorded for an additional 30 min. ΔFRET represents the FRET signal in the presence of agonist, minus the FRET signal over time in the presence of vehicle. Data are shown as mean±SEM from independent experiments as the mean value from triplicate wells.

#### Patch clamp recording

DRG (T11 - L3) were incubated with collagenase IV (1 mg/mL, Worthington) and dispase (4 mg/mL, Roche) (10 min, 37°C), were washed and dissociated with a fire-polished Pasteur pipette. Neurons were plated onto laminin- (0.017 mg/mL) and poly-D-Lysine- (2 mg/mL) -coated glass coverslips. Cells were incubated overnight in F12 medium containing 10% of fetal calf serum, 1% penicillin and streptomycin (Sigma No. P4333) in and humidified chamber (95% air, 5% CO_2_, 16h, 37°C). Changes in excitability of small-diameter (<30 pF capacitance) DRG neurons with properties of nociceptors were quantified by measuring rheobase (minimum input current to elicit an action potential) by whole-cell perforated patch-clamp recordings using Amphotericin B (240 µg/mL) (10). Only neurons with resting membrane potential more negative than -40 mV were analyzed. The recording chamber was continuously perfused with external solution at 2 mL/min. External solution was (mM): 140 NaCl, 5 KCl, 10 HEPES, 10 D-glucose, 1 MgCl_2_, 2 CaCl_2_; pH to 7.4 with 3 M NaOH. Pipette solution was (mM): 110 K-gluconate, 30 KCl, 10 HEPES, 1 MgCl_2_, 2 CaCl_2_; pH 7.25 with 1 M KOH. Neurons were incubated for 2 h with PAMAM-Chol-AZ (0.3 µg/mL, 100 nM AZ3451), PAMAM-Chol-Ø (0.3 µg/mL), PEG-Poly-AZ (0.1 µg/mL, 100 nM AZ3451), PEG-Poly-Chol-Ø (0.1 µg/mL) or unencapsulated AZ3451 (100 nM). Neurons were then washed, recovered for 90 min. incubated with trypsin (50 nM, 10 min), washed and rheobase was measured at T=30 min post-trypsin.

### Preclinical colitis models

#### 2F

PAMAM-Chol-Ø, (15 µg/mL), PAMAM-Chol-AZ3451 (∼15 µg/mL, 1, 5, 10 µM AZ3451), PEG-Poly-Ø (8 µg/mL), PEG-Poly-AZ (8 µg/mL, 5 µM AZ3451), unencapsulated AZ3451 (1, 5, 10 µM) (150 µl) or vehicle was injected into the colon by enema 3 cm from the anus of sedated mice (3-5% isoflurane). After 30 min, PAR_2_ agonist 2-furoyl-LIGRLO-NH_2_ (2F) (100 µM) or vehicle (80% NaCl, 10% Tween 20, 10% EtOH, control) (both 150 µl) was similarly administered. AZ3451 (5 µM, 200 µl) or vehicle was injected into the tail vein 30 min before intracolonic 2F. Withdrawal responses to abdominal stimulation with calibrated von Frey filaments (VFF) were measured for 6 h.

#### TNBS

2,4,6-Trinitrobenzene sulphonic acid (TNBS, Millipore Sigma, #P2297, 4 mg/mL) or vehicle (0.9% NaCl, 50% ethanol) (150 µl) was injected into the colon as described above (1). Body weight was monitored daily. After 96 h, PAMAM-Chol-Ø, 15 µg/mL), PAMAM-Chol-AZ3451 (15 µg/mL, 5 µM AZ3451), PEG-Poly-Ø (8 µg/mL), PEG-Poly-AZ (8 µg/mL, 5 µM AZ3451), unencapsulated AZ3451 (5 µM) (150 µl) or vehicle was similarly administered. AZ3451 (5 µM, 200 µl) or vehicle was injected into the tail vein 96 h after TNBS. Withdrawal responses to abdominal stimulation with calibrated VFF was measured for 6 h. The spleen and colon were removed. Spleen weight and colon length and cytokine and chemokine expression were measured.

#### DSS

Dextran sodium sulphate (DSS, MP Biomedicals, #0216011050, 3%) was administered in drinking water for 7 days; controls received plain water (1). Body weight, stool consistency and signs of rectal bleeding were recorded daily. The disease activity index (DAI) was determined by scoring body weight loss (0, none; 1, 1-5%; 2, 5-10%; 3, 10-20%; 4, >20%), stool consistency (0, normal; 2, loose stools; 4, diarrhea), and stool blood (0, negative; 2, positive hemoccult test; 4, gross bleeding). On day 8, PAMAM-Chol-Ø, 15 µg/mL), PAMAM-Chol-AZ3451 (15 µg/mL, 5 µM AZ3451), PEG-Poly-Ø (8 µg/mL), PEG-Poly-AZ (8 µg/mL, 5 µM AZ3451), unencapsulated AZ3451 (5 µM) (150 µl) or vehicle was similarly administered. Withdrawal responses to abdominal stimulation with calibrated VFF were measured for 6 h. The colon was removed and length and cytokine and chemokine expression were measured. Spleen weight was determined.

### Colonic nociception

The abdominal area was divided into 9 equal quadrants; von Frey filaments (VFF) of increasing force (0.07-2 g) were applied to the central quadrant, which corresponds to the area of the colon (1). Responses to abdominal stimulation were arching the back, jumping and raising/kicking the rear legs. Responses were measured hourly for at least 6 h. Results are expressed as VFF withdrawal threshold (g). Investigators were blinded to treatments.

### Spontaneous non-evoked behavior

Spontaneous behavior was assessed using a spectrometer (Behavior Sequencer, Behavioral Instruments, NJ; BiObserve) that objectively quantifies locomotor, exploratory and grooming behavior (11, 12). The spectrometer is a 40 cm^2^ arena with a CCD camera mounted in the center of the ceiling and a door aperture in the front area of the arena. Mouse movement was assessed by a floor mounted vibration sensor and 32 wall mounted infrared transmitter and receiver pairs. Mice were individually placed in the center of the behavioral spectrometer and their behavior was recorded, tracked, evaluated, and analyzed using a computerized video tracking system (Viewer3, BiObserve, DE) for 30 min. Total distance traveled in the open field, average velocity of locomotion, wall distance, ambulation and grooming were recorded and analyzed.

### Colonic inflammation

RNA was isolated from snap-frozen tissues using a Direct-zol RNA Kit (Zymo Research, #R2050). cDNA was synthetized from 50 ng of DNase-treated RNA using a High-Capacity cDNA Reverse Transcriptase Kit (Applied Biosystems, #4374966,). cDNA (50 or 100 ng) was amplified for 40 cycles by qRT-PCR using the QuantStudio 3 Real-Time PCR System. TNFα (#Mm00443258_m1), IL-1β (#Mm00434228_m1) and CXCL1 (#Mm04207460_m1) mRNA expression was measured by qPCR using TaqMan® gene expression and TaqMan® Fast Advanced Master Mix (#4444557). Relative amounts of target genes were calculated by normalization to the expression of with GAPDH reference gene (#Mm9999915_g1). Reference and target genes level were compared using the 2-ΔΔCT method (1).

### Data analysis and statistics

Sample size was pre-determined by past experience and by a power analysis of our previous studies. The sample size was not altered after the study commenced. Observations of cultured cell lines were replicated in n>5 independent experiments, with triplicate measurements in each experiment. Studies of cultured neurons were replicated in >12 neurons isolated from >4 mice. Experiments in mice were replicated in n>5 animals. The rules for stopping data collection were established in advance. Studies of cells were ended at predetermined times. Studies of mice with colitis were terminated if body weight was reduced by >20% during the experimental period. Inclusion and exclusion criteria for studies of cells and animal were established in advance. Inclusion criteria for cells included confluency. Inclusion criteria for mice included health status, age, sex and weight/body condition. Exclusion criteria for cells included contamination. Exclusion criteria for animal studies included wight loss >20%. All collected data were included in the study. No data were excluded. Outliers were not excluded from the study. Results were analyzed and graphs prepared using Prism 10. Data sets were normality tested with a D’Agostino & Pearson and Shapiro-Wilk test and analyzed statistical significance using one- or two-way ANOVA or Kruskal-Wallis test with Tuckey’s, Šidák’s, Bonferroni’s or Dunn’s post hoc test. Results are expressed as mean±SEM. The level of significance was set at up *P* ≤ 0.05.

## Author Contributions

Conceptualization: NMP, KL, BLS, NWB. Methodology: LTP, MB. Investigation: ST, RL, DB, PL, RP, CJP, B.S., GSAT, TC, DDJ, NNJ, AM. Visualization: RL, ST, DB, CJP. Supervision: SJV, NMP, KL, BLS, NWB. Funding acquisition: MB, NMP, KL, BLS, NWB. Writing – original draft: ST, RL, DB, CJP, BS, NNJ. Writing – review & editing: NMP, BLS, NWB.

## Classification

Biological Science. Pharmacology

## SI Figures

**Figure S1.**
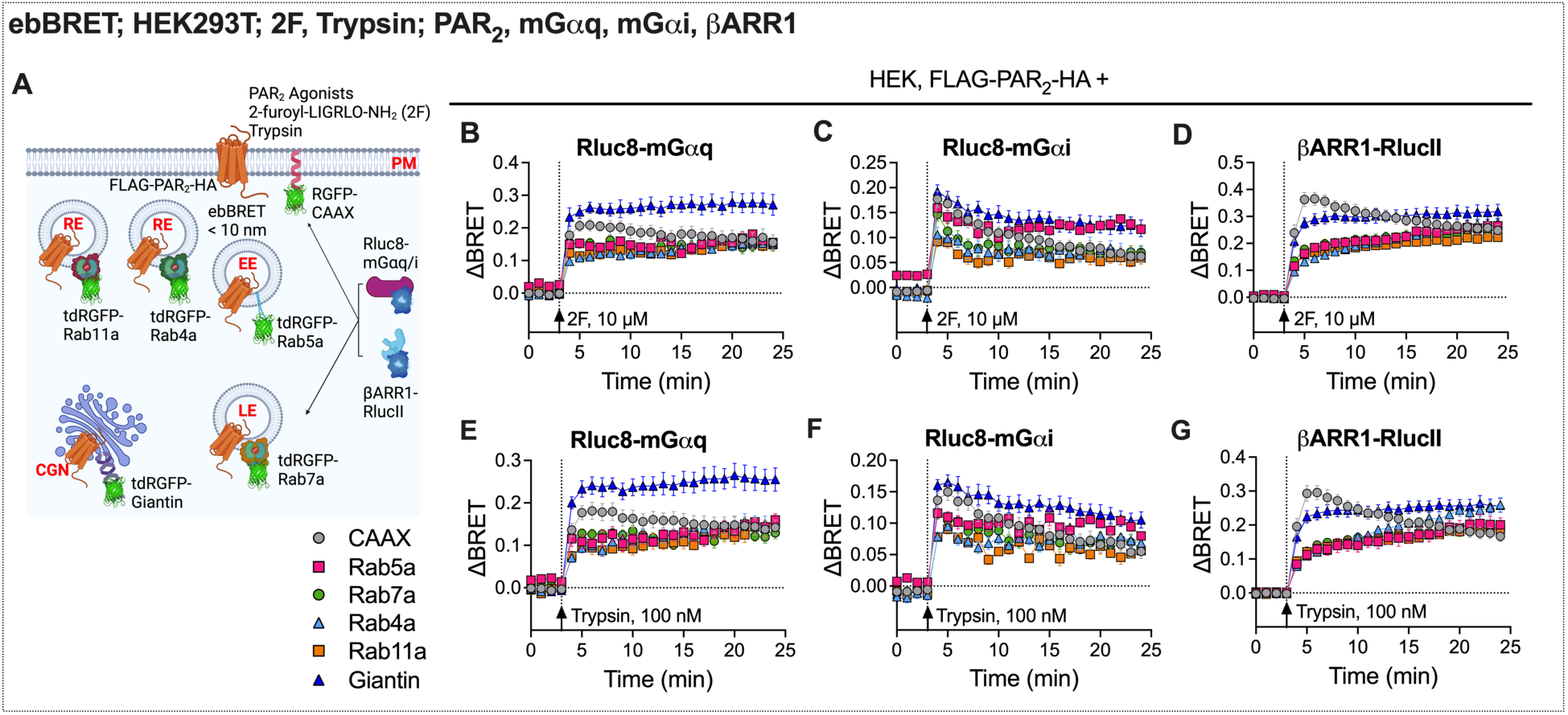
PAR_2_ agonists stimulate recruitment of mGα and βARR1 to the plasma membrane, endosomes and the Golgi apparatus. **A**. EbBRET assays of 2F-stimulated (**B-D**) or trypsin-stimulated (**E-G**) recruitment of mGαq (**B,E**), mGαi (**C,F**), and βARR1 (**D,G**) to the plasma membrane (CAAX), early (Rab5a), late (Rab7a), recycling (Rab4a and Rab11a) endosomes, and *cis-*Golgi network (Giantin) in HEK293T cells. Mean±SEM, triplicate observations from n=4-5 independent experiments.

**Figure S2.**
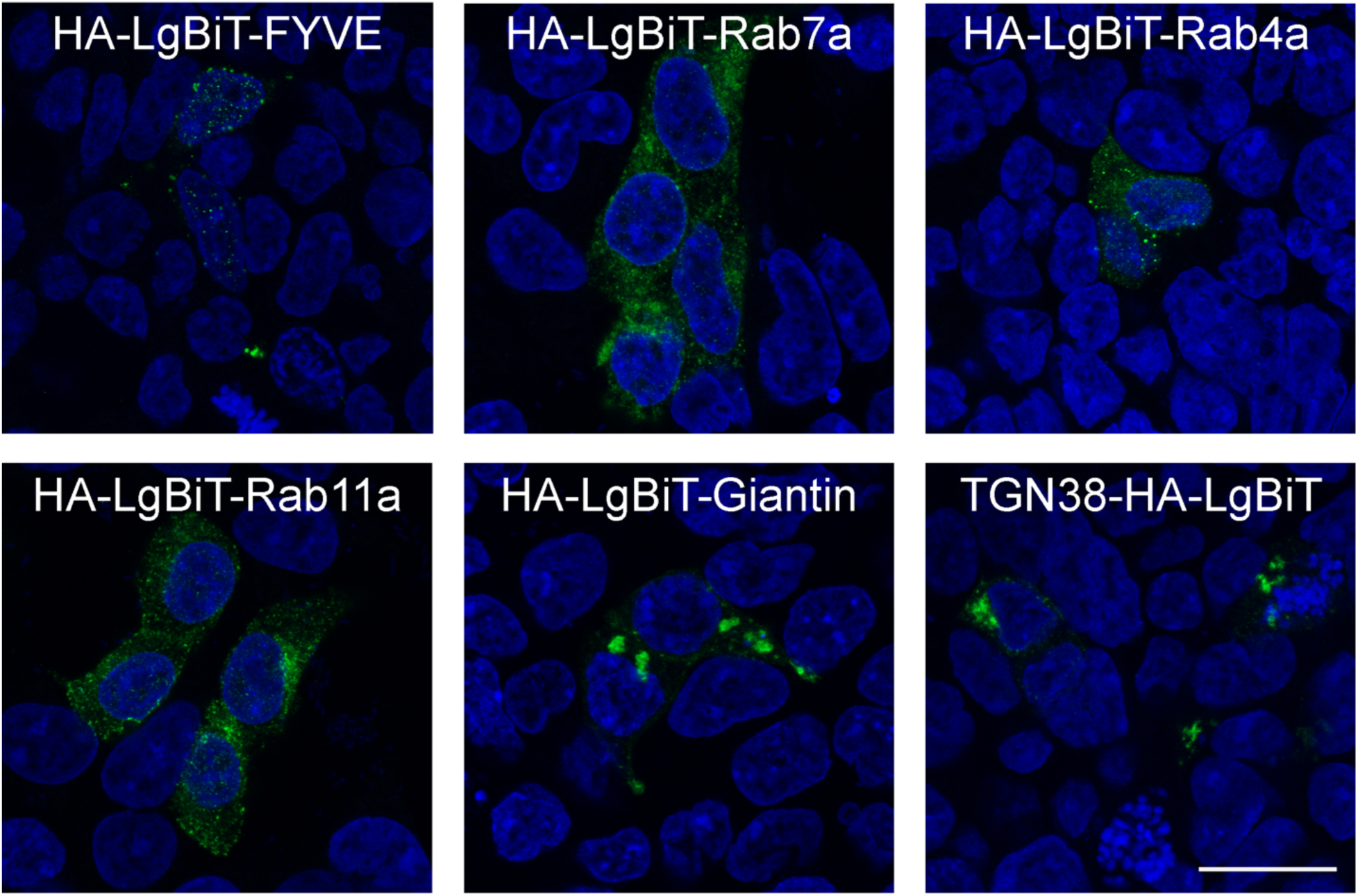
Localization markers for nbBRET assays are targeted to subcellular compartments. Confocal images showing localization of nbBRET markers of early endosomes (FYVE), late endosomes (Rab7a), recycling endosomes (Rab4a, Rab11a), *cis*-Golgi (Giantin) and *trans*-Golgi (TGN38). Markers were detected by immunofluorescence staining for HA11. Representative images, n=5 independent experiments. Scale bar, 20 µm.

**Figure S3.**
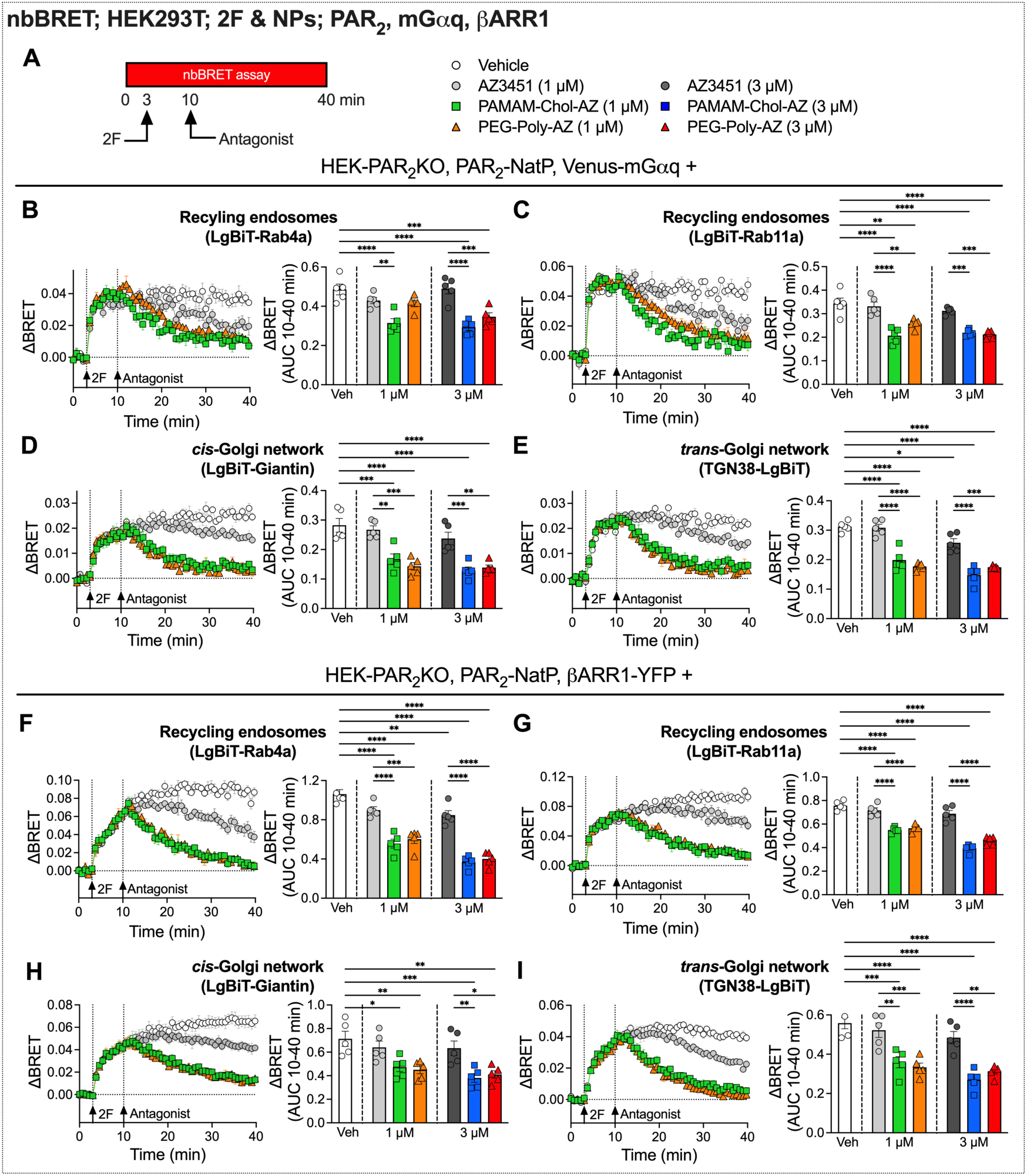
NPs encapsulating a PAR_2_ antagonist reverse 2F-stimulated PAR_2_ and βARR signaling in recycling endosomes and the Golgi apparatus. **A**. NbBRET assays of PAR_2_ antagonist reversal of 2F-stimulated PAR_2_ and mGαq/βARR1 compartmentalized signaling in HEK293T cells. **B-E**. 2F-induced recruitment of mGαq to PAR_2_ in recycling endosomes (Rab4a, Rab11a), *cis*-Golgi network (Giantin), and *trans*-Golgi network (TGN38) followed by treatment with vehicle, unencapsulated AZ3451, PAMAM-Chol-AZ, or PEG-Poly-AZ (1 µM or 3 µM). Time courses show responses to 2F followed by vehicle or antagonist (1 µM). Area under curve (AUC, 10-40 min) show integrated responses to 2F after vehicle or antagonist (1 or 3 µM). **F-I.** 2F-induced recruitment of βARR1 to PAR_2_ in recycling endosomes (Rab4a, Rab11a), *cis*-Golgi network (Giantin), and *trans*-Golgi network (TGN38) followed by treatment of vehicle, unencapsulated AZ3451, PAMAM-Chol-AZ, or PEG-Poly-AZ (1 µM or 3 µM). Time courses show responses to 2F followed by vehicle or antagonist (1 µM). Area under curve (AUC, 10-40 min) show integrated responses to 2F after vehicle or antagonist (1 or 3 µM). Mean±SEM, triplicate observations, n=5 independent experiments. \**P*<0.05, \*\**P*<0.01, \*\*\**P*<0.001, \*\*\*\**P*<0.0001. One-way ANOVA, Šidák’s test.

**Figure S4.**
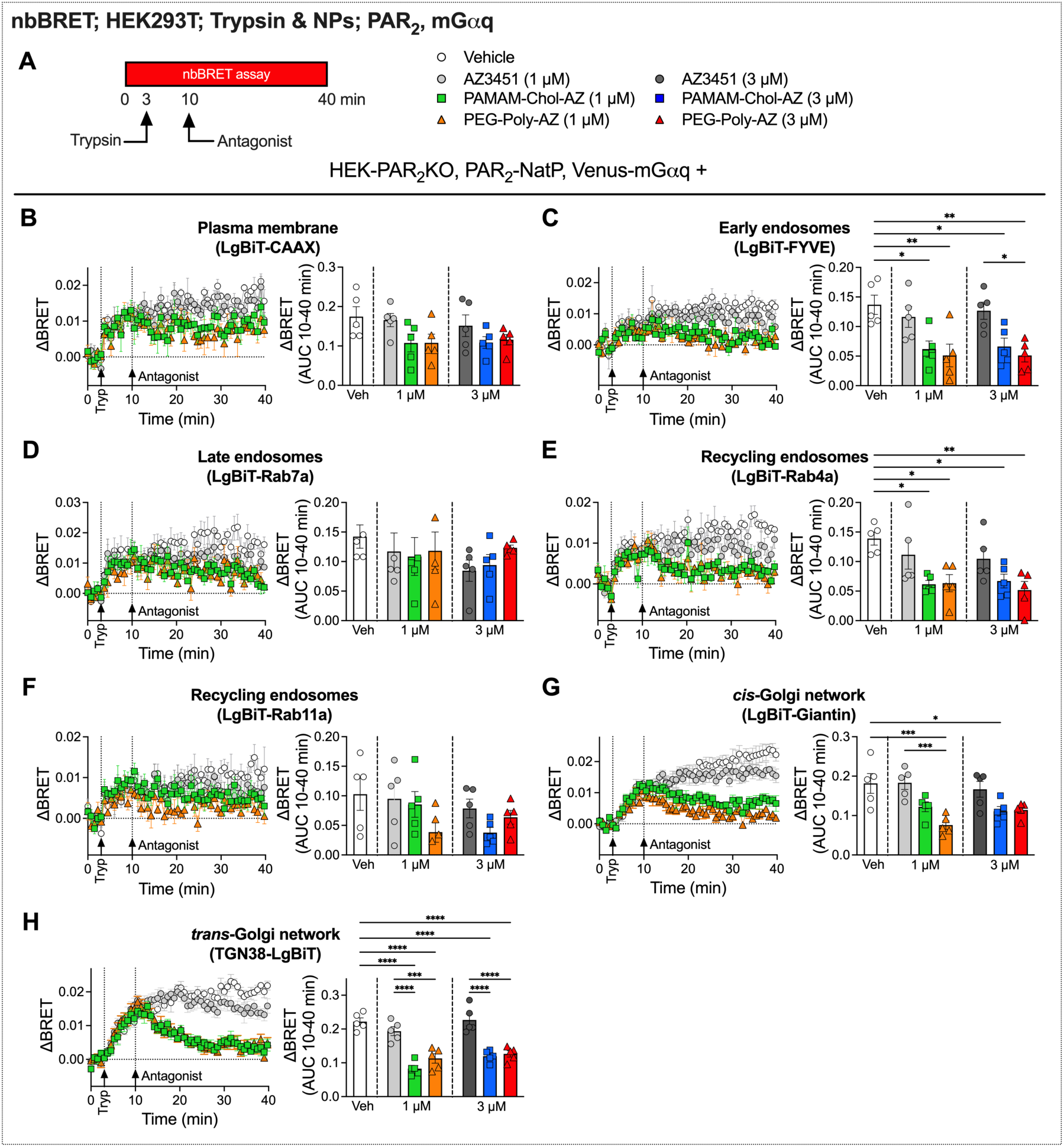
NPs encapsulating a PAR_2_ antagonist reverse trypsin-stimulated PAR_2_/mGαq signaling in endosomes and the Golgi apparatus. **A.** NbBRET assays of PAR_2_ antagonist reversal of trypsin-stimulated PAR_2_ and mGαq compartmentalized signaling in HEK293T cells. **B-H**. Trypsin-induced recruitment of mGαq to PAR_2_ to the plasma membrane (CAAX), early endosomes (FYVE), late endosomes (Rab7a), recycling endosomes (Rab4a, Rab11a), *cis*-Golgi network (Giantin), and *trans*-Golgi network (TGN38) followed by treatment of vehicle, unencapsulated AZ3451, PAMAM-Chol-AZ, or PEG-Poly-AZ (1 µM or 3 µM). Time courses show responses to trypsin followed by vehicle or antagonist (1 µM). Area under curve (AUC, 10-40 min) show integrated responses to trypsin after vehicle or antagonist (1 or 3 µM). Mean±SEM, triplicate observations, n=5 independent experiments. \**P*<0.05, \*\**P*<0.01, \*\*\**P*<0.001, \*\*\*\**P*<0.0001. One-way ANOVA, Šidák’s test.

**Figure S5.**
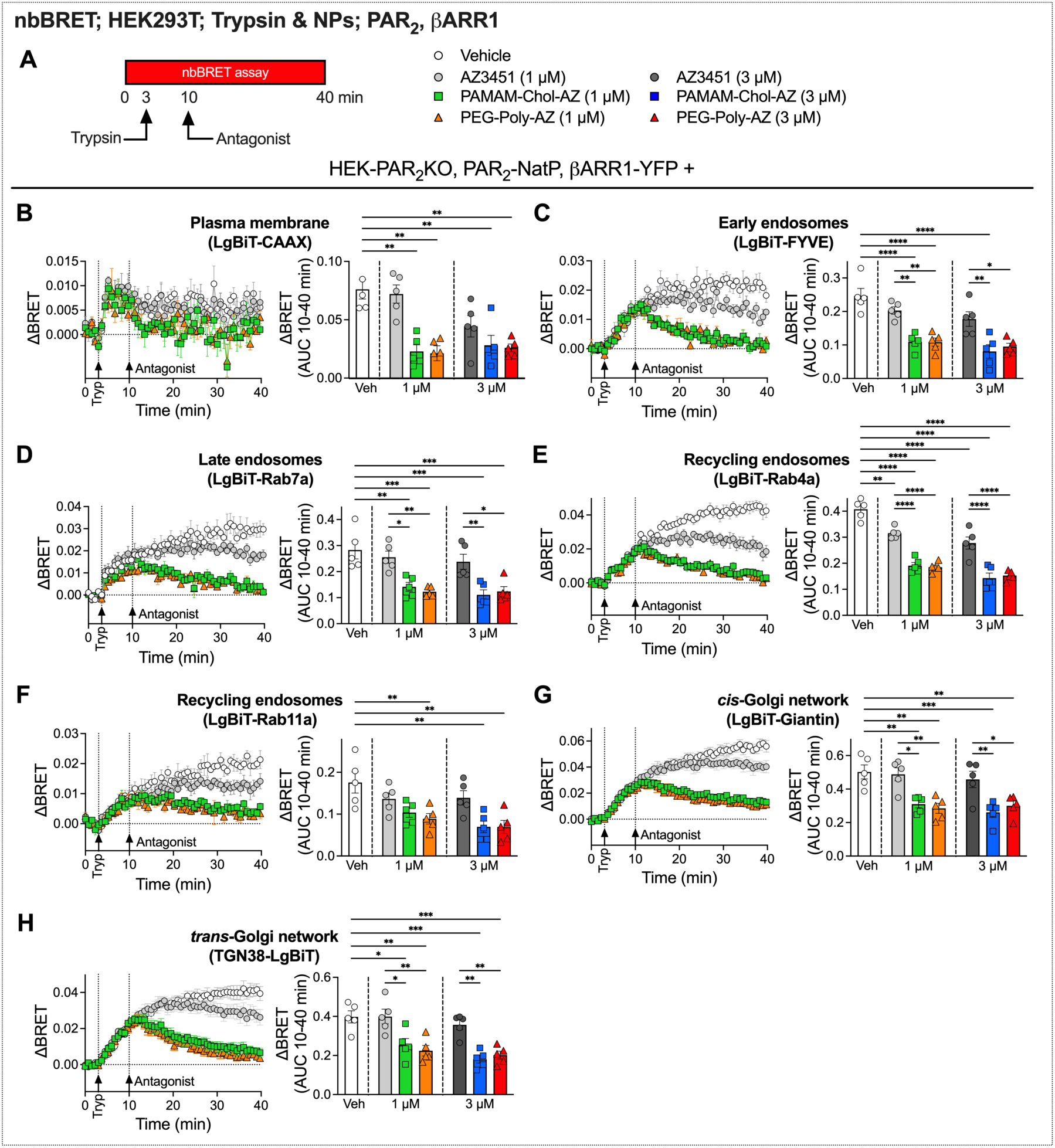
NPs encapsulating a PAR_2_ antagonist reverse trypsin-stimulated PAR_2_/βARR1 signaling in endosomes and the Golgi apparatus. **A.** NbBRET assays of PAR_2_ antagonist reversal of trypsin-stimulated PAR_2_/βARR1 compartmentalized signaling in HEK293T cells. **B-H**. Trypsin-induced recruitment of βARR1 to PAR_2_ at the plasma membrane (CAAX), early endosomes (FYVE), late endosomes (Rab7a), recycling endosomes (Rab4a, Rab11a), *cis*-Golgi network (Giantin), and *trans*-Golgi network (TGN38) followed by treatment of vehicle, unencapsulated AZ3451, PAMAM-Chol-AZ, or PEG-Poly-AZ (1 µM or 3 µM). Time courses show responses to trypsin followed by vehicle or antagonist (1 µM). Area under curve (AUC, 10-40 min) show integrated responses to trypsin after vehicle or antagonist (1 or 3 µM). Mean±SEM, triplicate observations, n=5 independent experiments. \**P*<0.05, \*\**P*<0.01, \*\*\**P*<0.001, \*\*\*\**P*<0.0001. One-way ANOVA, Šidák’s test.

**Figure S6.**
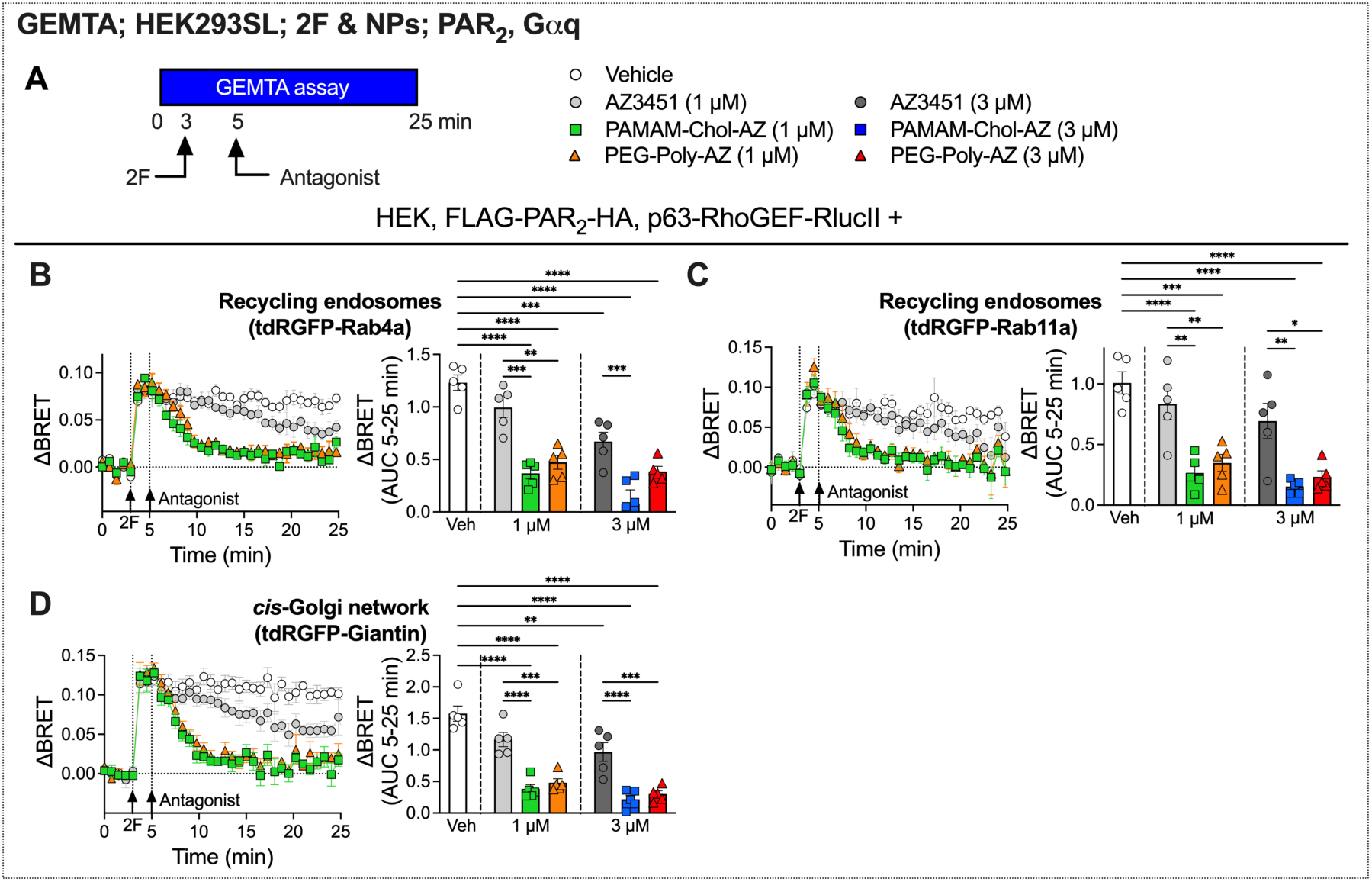
NPs encapsulating a PAR_2_ antagonist reverse 2F-stimulated Gαq signaling in recycling endosomes and the Golgi apparatus. **A**. GEMTA assays of PAR_2_ antagonist reversal of 2F-stimulated PAR_2_ and Gαq compartmentalized signaling in HEK293T cells. **B-D**. 2F-induced recruitment of p63-RhoGEF to recycling endosomes (Rab4a, Rab11a) and *cis*-Golgi (Giantin) followed by treatment with vehicle, unencapsulated AZ3451, PAMAM-Chol-AZ, or PEG-Poly-AZ (1 µM or 3 µM). Time courses show responses to 2F followed by vehicle or antagonist (1 µM). Area under curve (AUC, 5-25 min) show integrated responses to 2F after vehicle or antagonist (1 or 3 µM). Mean±SEM, triplicate observations, n=5 experiments. \**P*<0.05, \*\**P*<0.01, \*\*\**P*<0.001, \*\*\*\**P*<0.0001. One-way ANOVA, Šidák’s test.

**Figure S7.**
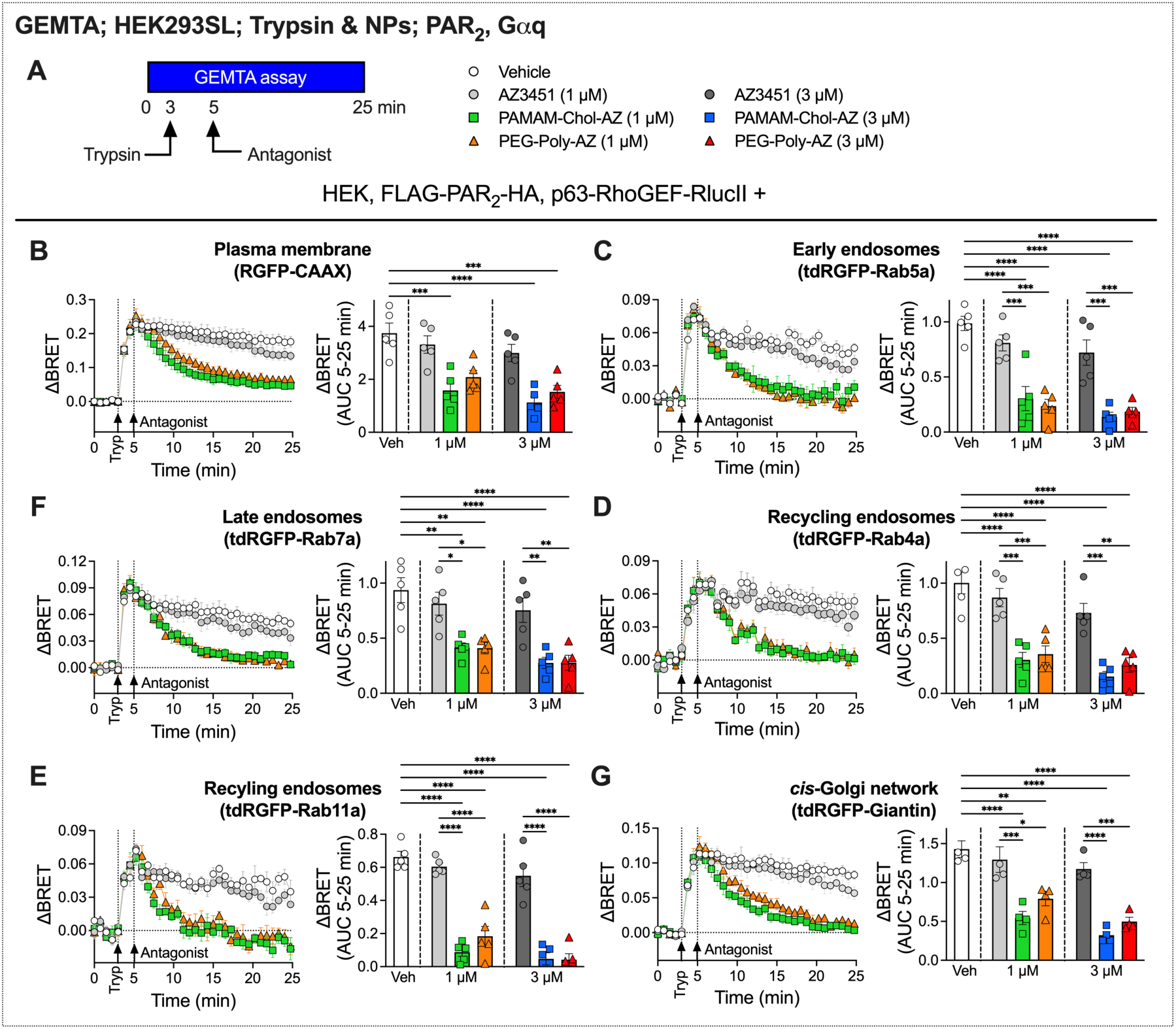
NPs encapsulating a PAR_2_ antagonist reverse trypsin-stimulated Gαq signaling in endosomes and the Golgi apparatus. **A**. GEMTA assays of PAR_2_ antagonist reversal of trypsin-stimulated PAR_2_ and Gαq compartmentalized signaling in HEK293T cells. **B-G**. Trypsin-induced recruitment of p63-RhoGEF to the plasma membrane (CAAX), early endosomes (Rab5a), late endosomes (Rab7a), recycling endosomes (Rab4a, Rab11a) and *cis*-Golgi network (Giantin) followed by treatment with vehicle, unencapsulated AZ3451, PAMAM-Chol-AZ, or PEG-Poly-AZ (1 µM or 3 µM). Time courses show responses to trypsin followed by vehicle or antagonist (1 µM). Area under curve (AUC, 5-25 min) show integrated responses to trypsin after vehicle or antagonist (1 or 3 µM). Mean±SEM, triplicate observations, n=5 experiments. \**P*<0.05, \*\**P*<0.01, \*\*\**P*<0.001, \*\*\*\**P*<0.0001. One-way ANOVA, Šidák’s test.

**Figure S8.**
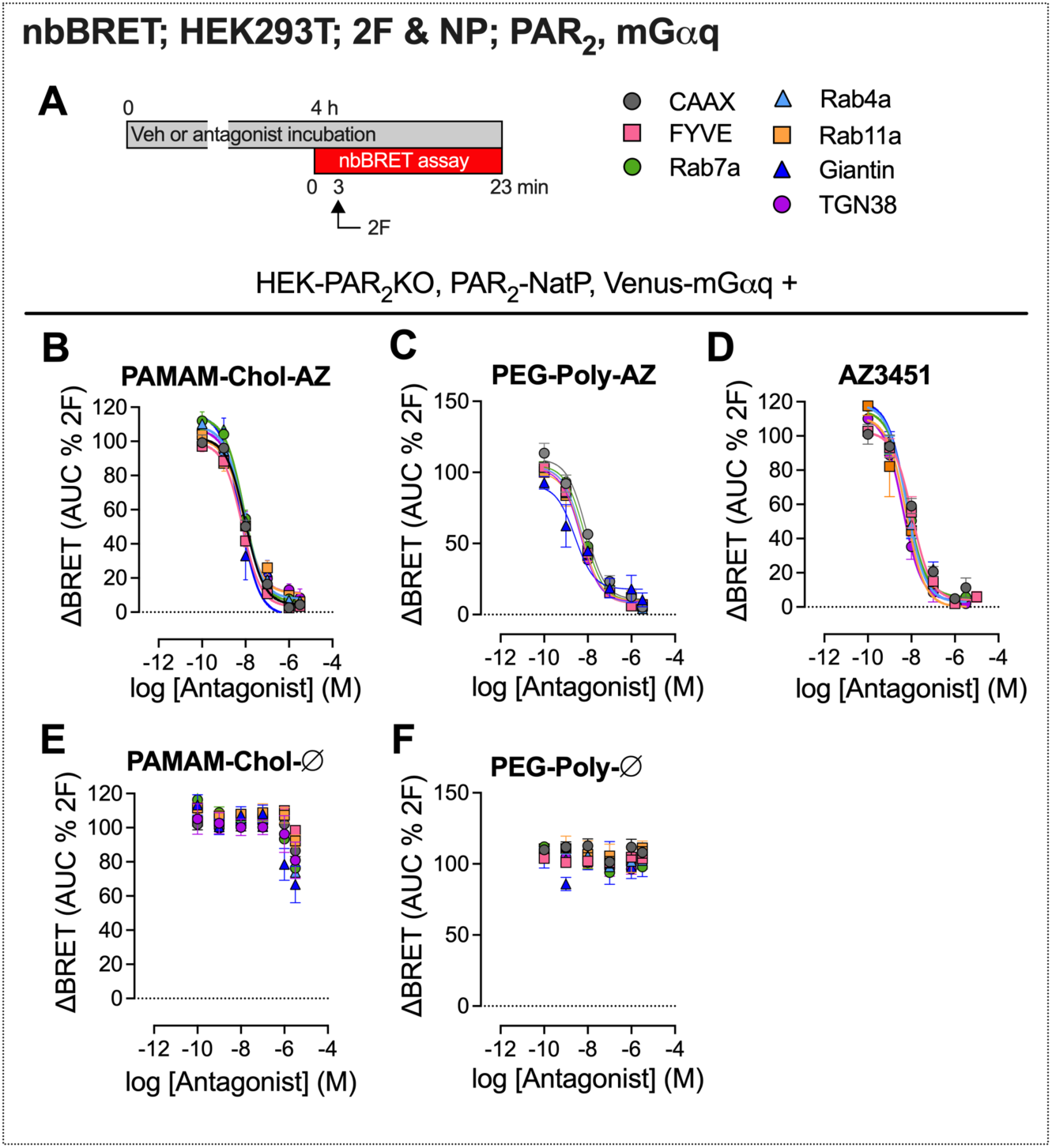
NP-encapsulated AZ3451 retains full PAR_2_ antagonist activity. **A**. NbBRET assays of effects of preincubation with NPs encapsulated with AZ3451 on PAR_2_ and mGαq compartmentalized signaling in HEK293T cells. **B-F**. Concentration-response to 2F-induced recruitment of mGαq to PAR_2_ at the plasma membrane (CAAX), early endosomes (FYVE), late endosomes (Rab7a), recycling endosomes (Rab4a, Rab11a), *cis*-Golgi network (Giantin), or *trans*-Golgi network (TGN38) of HEK293T cells preincubated for 4 h with vehicle, graded concentrations of PAMAM-Chol-AZ (**B**), PEG-Poly-AZ (**C**), unencapsulated AZ3451 (**D**), PAMAM-Chol-Ø (**E**) and PAMAM-Chol-Ø (**F**). Mean±SEM, triplicate observations from n=5 independent experiments.

**Figure S9.**
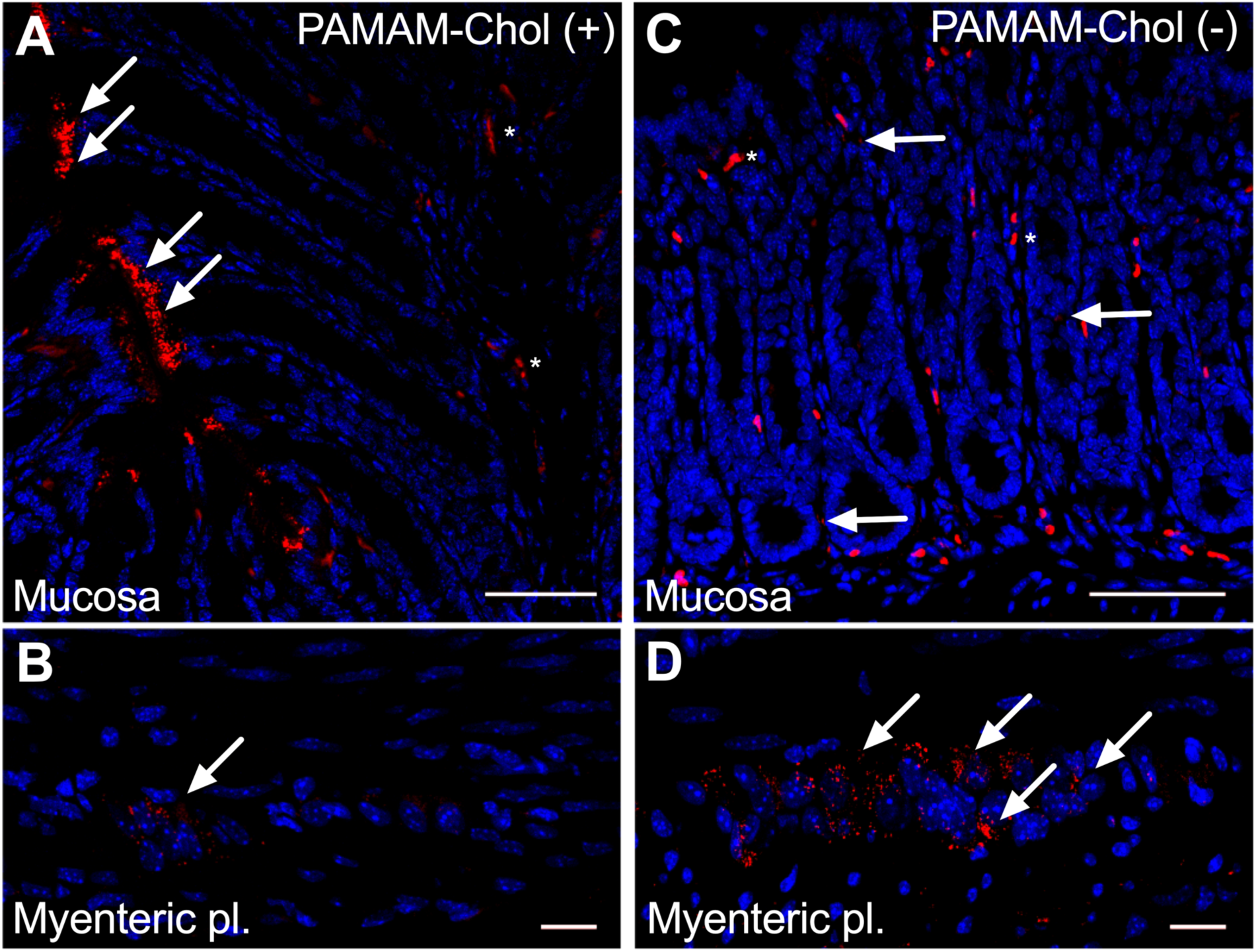
Surface charge determines cellular targeting of NPs in the colon. **A-D.** Localization of PAMAM-Chol-Cy5(+) (**A, B**) and PAMAM-Chol-Cy5(-) (**C, D**) in the mucosa and myenteric plexus (pl.) of the colon at 2 h after intraluminal injection. Arrows denote PAMAM-Chol-Cy5 NPs and * denotes auto-fluorescent cells in the mucosa. Representative images from experiments in n=3 mice. Scale, 20 µm.

**Figure S10.**
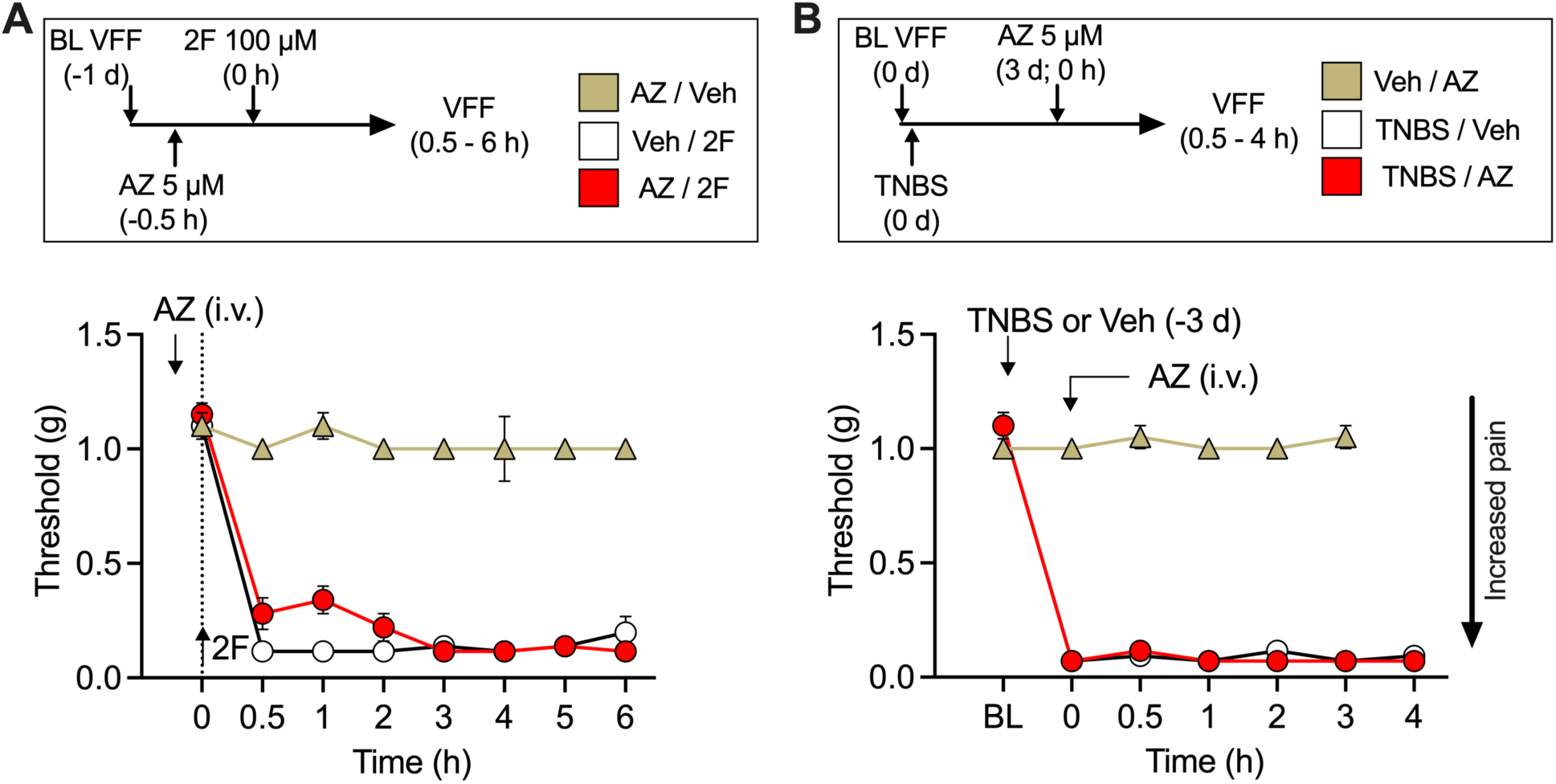
Systemic PAR_2_ antagonist AZ3451 does not affect colonic nociception. **A.** AZ3451 or vehicle (Veh) was injected intravenously (i.v.) 30 min before intracolonic injection of 2F. **B.** AZ3451 was injected intravenously 3 d after intracolonic injection of TNBS. Abdominal withdrawal threshold to VFF stimulation was measured for up to 6 h after AZ3451. Mean±SEM, n=5 mice.

**Figure S11.**
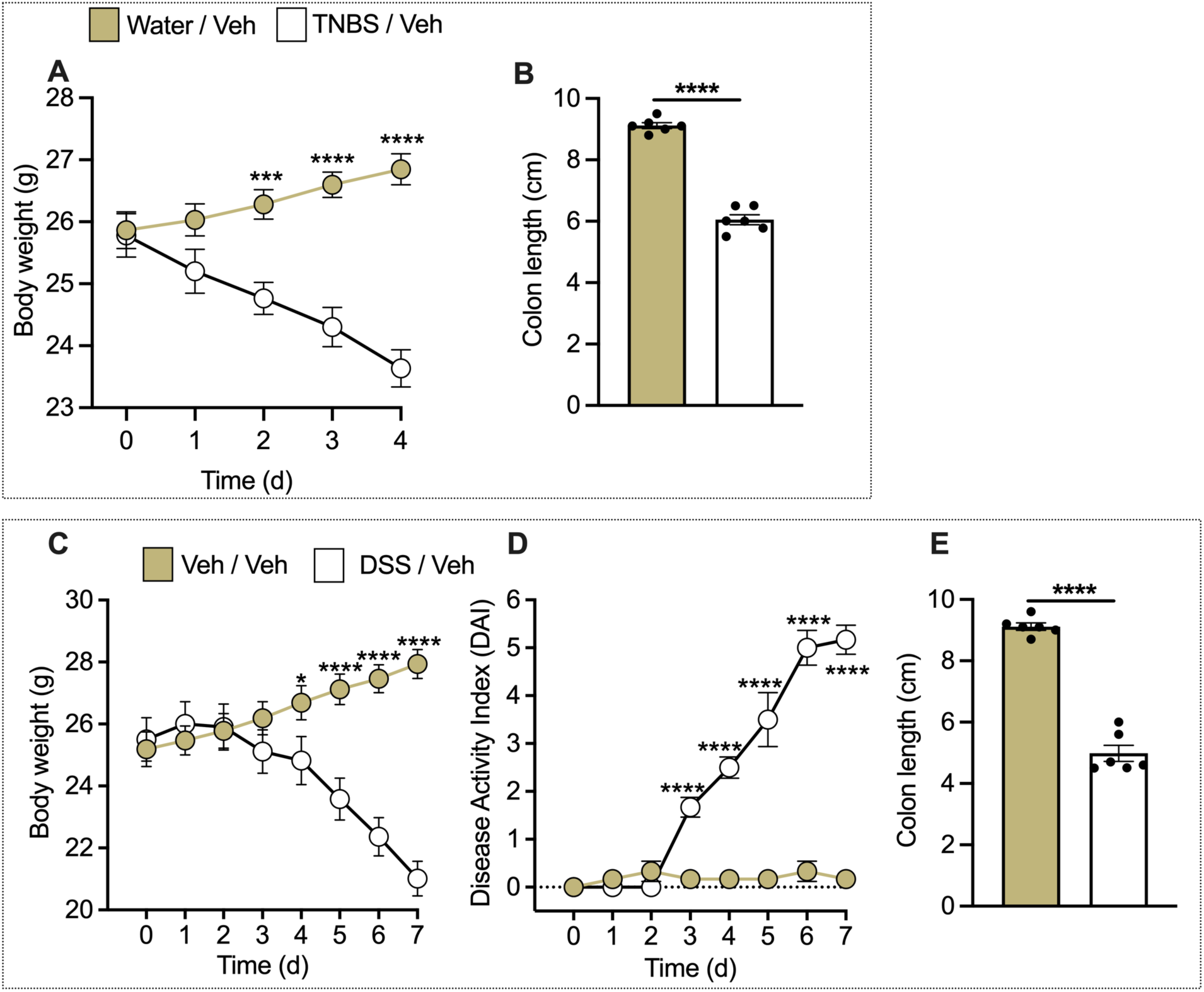
TNBS and DSS induce signs of colonic inflammation in mice. **A, B.** TNBS colitis. **C-E.** DSS colitis. **A, C.** Change in body weight. **D.** Disease activity index. **B, E.** Colon length at day 4 (TNBS) and day 7 (DSS). Mean±SEM, n=6 mice. \*\*\**P*<0.001, \*\*\*\**P*<0.0001. **A, C-D.** Two-way ANOVA, Šidák’s multiple comparison test. **B, E.** Unpaired t-test.

**Figure S12.**
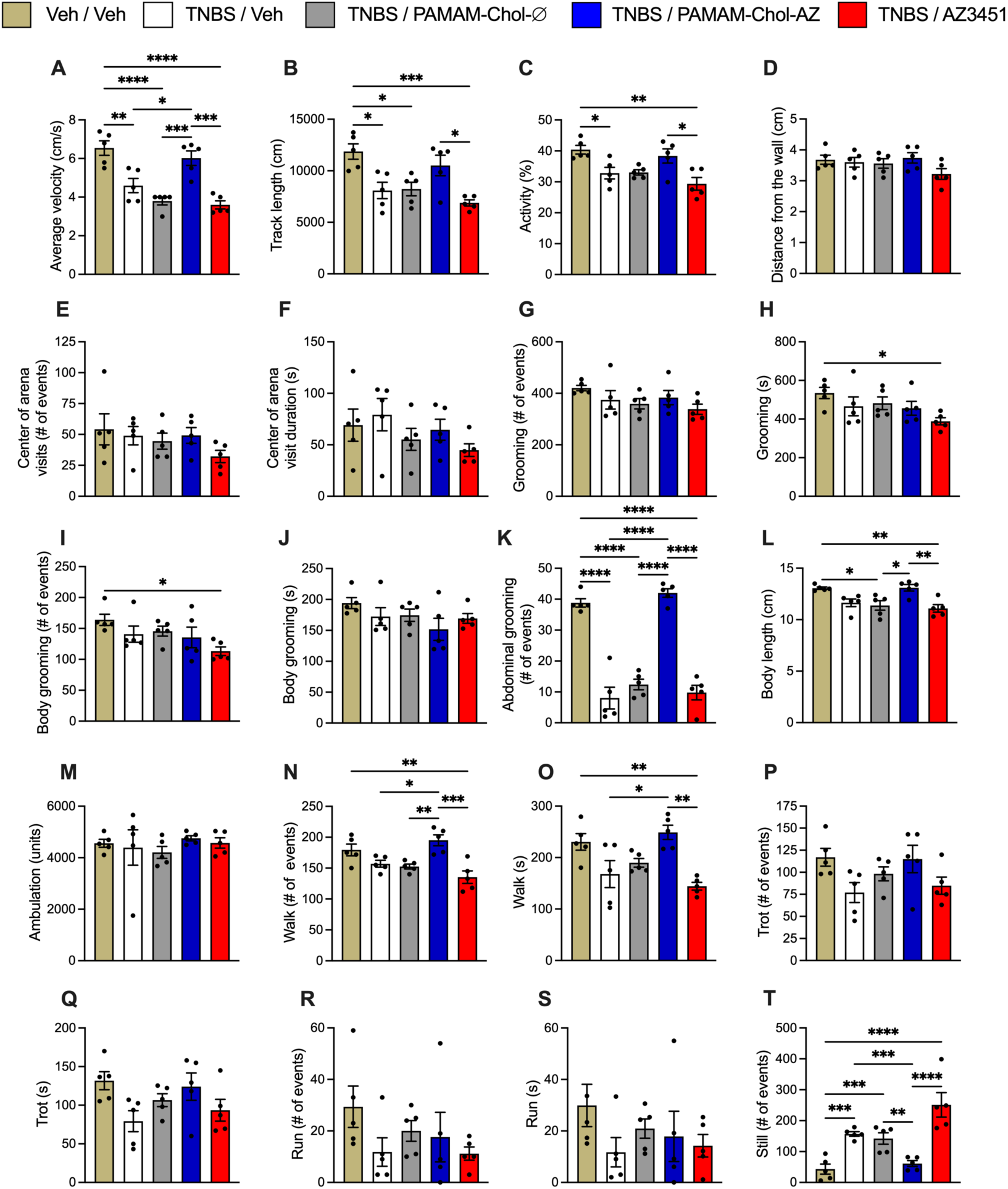
Luminally administered PAMAM-Chol-AZ NPs normalize aberrant behavior in mice with TNBS colitis. At 3 d after vehicle or TNBS treatment, vehicle, PAMAM-Chol-Ø, PAMAM-Chol-AZ or unencapsulated AZ3451 was administered into the colon lumen. After 3 h, behavior was monitored for 30 min using a behavioral spectrometer. **A.** Average velocity. **B.** Track length. **C.** Activity. **D.** Distance from wall. **E.** Number of visits to center of arena. **F.** Duration of visits to the center of the arena. **G.** Grooming events. **H.** Grooming time. **I.** Body grooming events. **J.** Body grooming time. **K.** Abdominal grooming events. **L.** Body length. **M.** Ambulation. **N.** Walk events. **O.** Walk time. **P.** Trot events. **Q.** Trot time. **R.** Run events. **S.** Run time. **T.** Still events. Mean±SEM, n=5 or 6 mice. *P*<0.05, \*\**P*<0.01, \*\*\**P*<0.001, \*\*\*\**P*<0.0001. One-way ANOVA, Tukey’s multiple comparison test.

**Figure S13.**
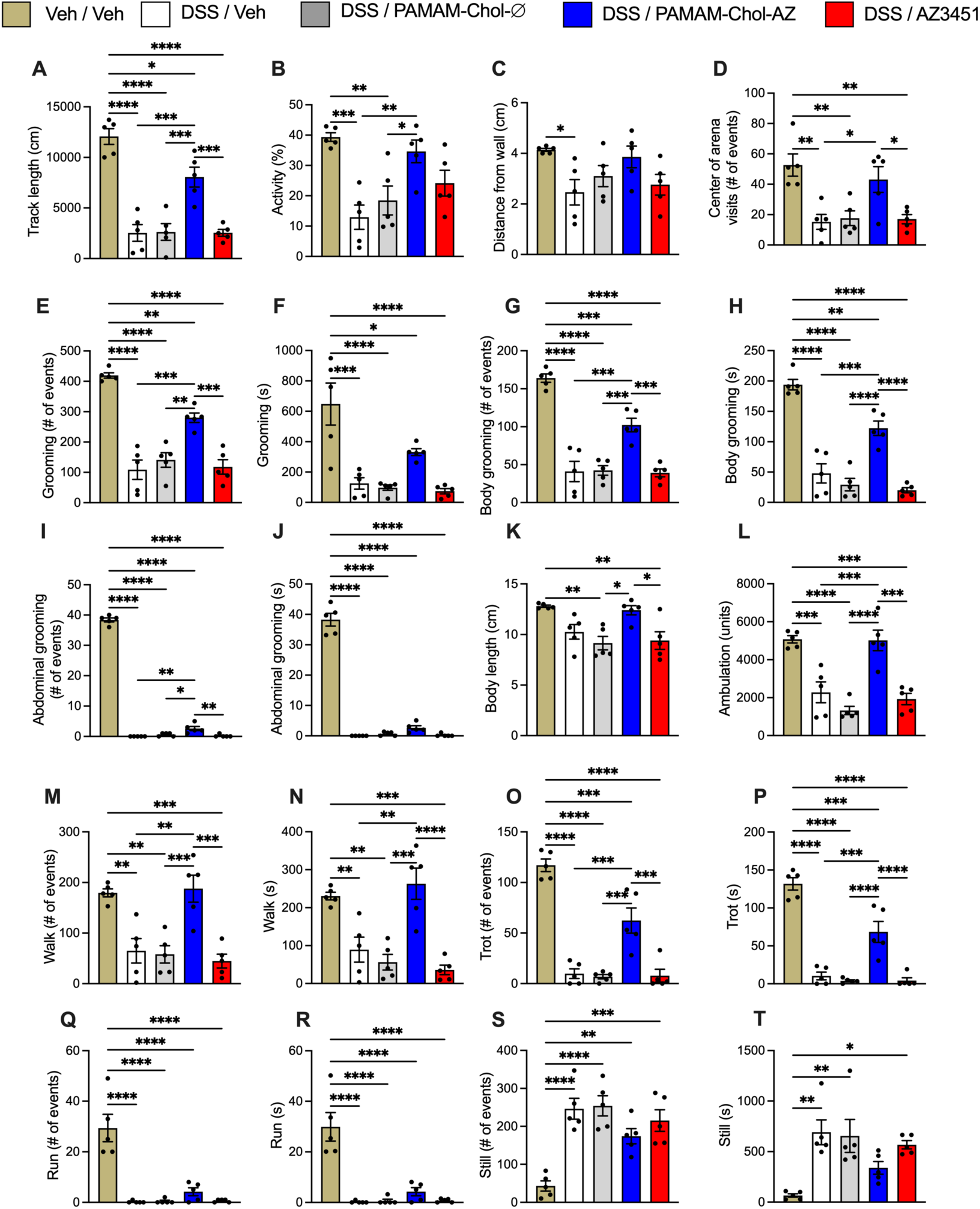
Luminally administered PAMAM-Chol-AZ NPs normalize aberrant behavior in mice with DSS colitis. At 8 d after starting water or DSS treatment, vehicle, PAMAM-Chol-Ø, PAMAM-Chol-AZ or unencapsulated AZ3451 was administered into the colon lumen. After 3 h, behavior was monitored for 30 min using a behavioral spectrometer. **A.** Track length. **B.** Activity. **C.** Distance from wall. **D.** Number of visits to the center of the arena. **E.** Grooming events. **F.** Grooming time. **G.** Body grooming events. **H.** Body grooming time. **I.** Abdominal grooming events. **J.** Abdominal grooming time. **K.** Body length. **L.** Ambulation. **M.** Walk events. **N.** Walk time. **O.** Trot events. **P.** Trot time. **Q.** Run events. **R.** Run time. **S.** Still events. **T.** Still time. Mean±SEM, n=5 or 6 mice. *P*<0.05, \*\**P*<0.01, \*\*\**P*<0.001, \*\*\*\**P*<0.0001. One-way ANOVA, Tukey’s multiple comparison test.

**Figure S14.**
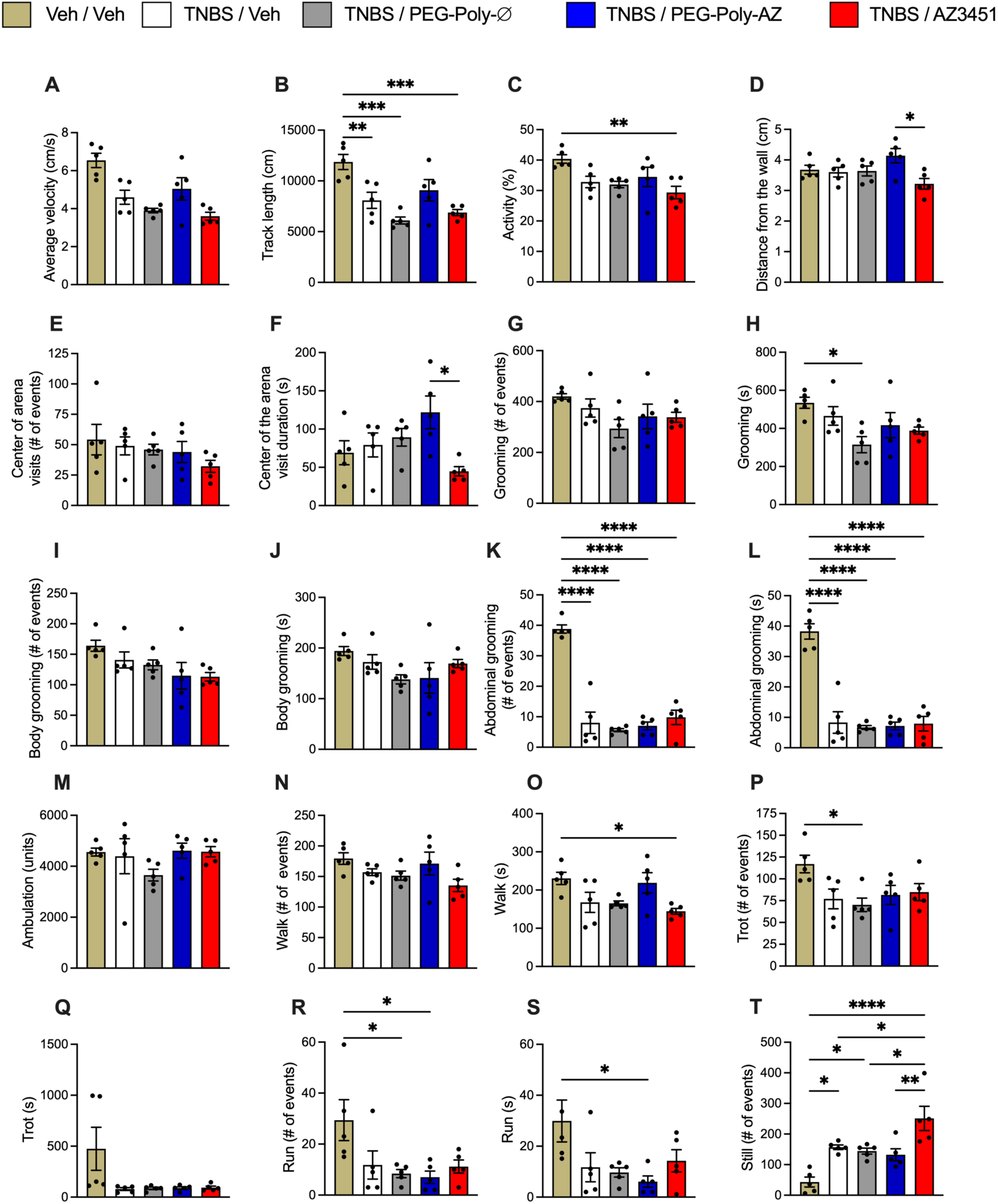
Luminally administered PEG-Poly-AZ NPs normalize aberrant behavior in mice with TNBS colitis. At 3 d after vehicle or TNBS treatment, vehicle, PEG-Poly-Ø, PEG-Poly-AZ or unencapsulated AZ3451 was administered into the colon lumen. After 3 h, behavior was monitored for 30 min using a behavioral spectrometer. **A.** Average velocity. **B.** Track length. **C.** Activity. **D.** Distance from wall. **E.** Number of visits to the center of the arena. **F.** Duration of visits to the center of the arena. **G.** Grooming events. **H.** Grooming time. **I.** Body grooming events. **J.** Body grooming time. **K.** Abdominal grooming events. **L.** Abdominal grooming time. **M.** Ambulation. **N.** Walk events. **O.** Walk time. **P.** Trot events. **Q.** Trot time. **R.** Run events. **S.** Run time. **T.** Still events. Mean±SEM, n=5 or 6 mice. *P*<0.05, \*\**P*<0.01, \*\*\**P*<0.001, \*\*\*\**P*<0.0001. One-way ANOVA, Tukey’s multiple comparison test.

**Figure S15.**
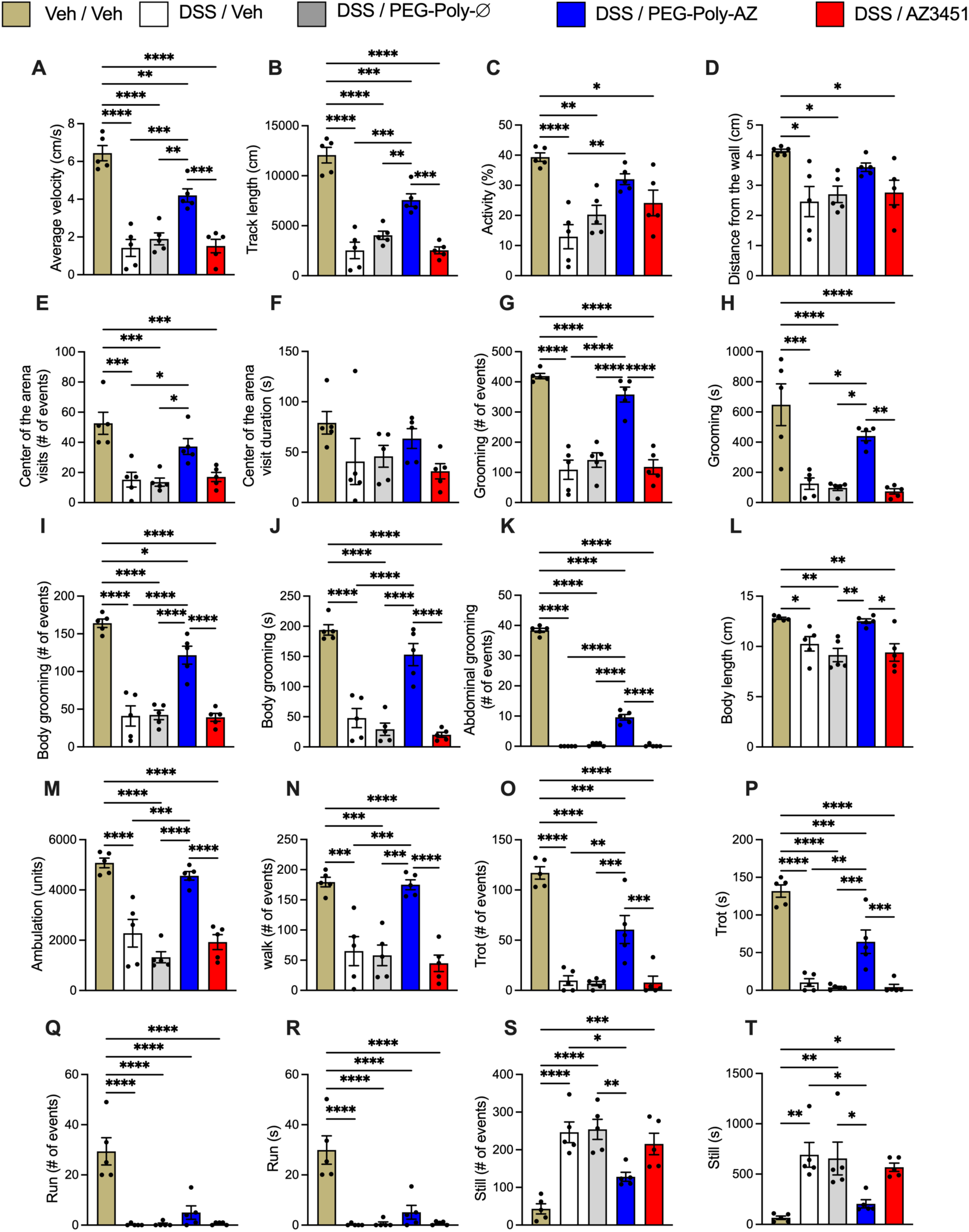
Luminally administered PEG-Poly-AZ NPs normalize aberrant behavior in mice with DSS colitis. At 8 d after starting water or DSS treatment, vehicle, PEG-Poly-Ø, PEG-Poly-AZ or unencapsulated AZ3451 was administered into the colon lumen. After 3 h, behavior was monitored for 30 min using a behavioral spectrometer. **A.** Average velocity. **B.** Track length. **C.** Activity. **D.** Distance from wall. **E.** Number of visits to center of arena. **F.** Duration of visits to arena. **G.** Grooming events. **H.** Grooming time. **I.** Body grooming events. **J.** Body grooming time. **K.** Abdominal grooming events. **L.** Body length. **M.** Ambulation. **N.** Walk events. **O.** Trot events. **P.** Trot time. **Q.** Run events. **R.** Run time. **S.** Still events. **T.** Still time. Mean±SEM, n=5 or 6 mice. *P*<0.05, \*\**P*<0.01, \*\*\**P*<0.001, \*\*\*\**P*<0.0001. One-way ANOVA, Tukey’s multiple comparison test.

## SI Movies

**Movie S1. Luminally administered PAMAM-Chol-AZ NPs normalize aberrant behavior of mice with TNBS colitis.** At 3 d after vehicle or TNBS treatment, vehicle or PAMAM-Chol-AZ was administered into the colon lumen. After 3 h, behavior was monitored for 30 min using a behavioral spectrometer. Representative recordings of n=6 mice per treatment. Recordings speeded up 3-fold.

**Movie S2. Luminally administered PAMAM-Chol-AZ NPs normalize aberrant behavior of mice with DSS colitis.** At 8 d after starting water or DSS treatment, vehicle or PAMAM-Chol-AZ was administered into the colon lumen. After 3 h, behavior was monitored for 30 min using a behavioral spectrometer. Representative recordings of n=6 mice per treatment. Recordings speeded up 3-fold.

**Movie S3. Luminally administered PEG-Poly-AZ NPs normalize aberrant behavior of mice with TNBS colitis.** At 3 d after vehicle or TNBS treatment, vehicle or PPEG-Poly-AZ was administered into the colon lumen. After 3 h, behavior was monitored for 30 min using a behavioral spectrometer. Representative recordings of n=6 mice per treatment. Recordings speeded up 3-fold.

**Movie S4. Luminally administered PEG-Poly-AZ NPs normalize aberrant behavior of mice with DSS colitis.** At 8 d after starting water or DSS treatment, vehicle or PEG-Polyl-AZ was administered into the colon lumen. After 3 h, behavior was monitored for 30 min using a behavioral spectrometer. Representative recordings of n=6 mice per treatment. Recordings speeded up 3-fold.

